# Mitotic interhomolog recombination drives genomic diversity in diatoms

**DOI:** 10.1101/2020.11.08.373134

**Authors:** Petra Bulánková, Mirna Sekulić, Denis Jallet, Charlotte Nef, Tom Delmont, Cock van Oosterhout, Ilse Vercauteren, Cristina Maria Osuna-Cruz, Emmelien Vancaester, Thomas Mock, Koen Sabbe, Fayza Daboussi, Chris Bowler, Wim Vyverman, Klaas Vandepoele, Lieven De Veylder

**Author notes:** These authors contributed equally to this work (Mirna Sekulić, Denis Jallet). Tree of Life, Wellcome Sanger Institute, Cambridge, CB10 1SA, UK.

## Abstract

Diatoms, an evolutionarily successful group of microalgae, display high levels of intraspecific variability in natural populations. However, the process generating such diversity is unknown. Here we estimated the variability within a natural diatom population and subsequently mapped the genomic changes arising within cultures clonally propagated from single diatom cells. We demonstrate that genome rearrangements and mitotic recombination between homologous chromosomes underlie clonal variability, resulting in haplotype diversity accompanied by the appearance of novel protein variants and loss of heterozygosity resulting in the fixation of alleles. The frequency of interhomolog mitotic recombination exceeds 4 out of 100 cell divisions and increases under environmental stress. We propose that this plastic response in the interhomolog mitotic recombination rate increases the evolutionary potential of diatoms, contributing to their ecological success.

**One Sentence Summary:** Recombination between homologous chromosomes in diatom vegetative cells leads to extensive genomic diversity in clonal populations.

## Main Text

Diatoms are an extraordinarily diverse and species-rich group, populating a wide range of aquatic environments from polar to tropical regions (*1*). Moreover, previous studies of intraspecific genetic variation in diatoms indicated high levels of variability in natural populations in species with differing life-cycle strategies (*2, 3*). While mutations are considered as a primary source of genetic variability, mutation rates in diatoms appear to be at a similarly low level as in green algae (*4*). Another source for genetic variability is sexual reproduction, typically generating new allelic combinations. However, several diatom species, including the model diatoms *Phaeodactylum tricornutum* and *Thalassiosira pseudonana*, have never been observed to produce F1 progeny (*5, 6*), whereas sex is restricted by cell size in many other diatoms (*7*). This opens a question: How can a group of organisms with an unexceptional mutation rate and only sporadic sexual reproduction generate such evolutionary novelty?

To first quantify the extent of intraspecific variability within a natural population we studied the intra-population variability of *Fragilariopsis cylindrus. F. cylindrus* is a marine diatom species with elevated genomic variability (*3*), and was highly detected in surface and deep chlorophyll maximum (DCM) zone metagenomes from *Tara* Oceans Station 86 in the Southern Ocean (near the Antarctic peninsula, 64°30’88” S, 53°05’75” W). A reference genome from an *F. cylindrus* culture genome was used to recruit metagenomic reads with high average nucleotide identity (ANI) to explore the intra-population variability. Single nucleotide variants (SNVs) distributed over 24,326 core genes, corresponding to 89.64% (30.95 Mb) of all *F. cylindrus* genes, resulted in 619,947 and 592,929 SNVs in the surface and DCM metagenomes, respectively, covering all the core genes (Data S1). This corresponds to a SNV density of ∼2%, corresponding to one SNV every 50 nucleotides. A-G and C-T transitions each contributed 30% of SNVs, followed by the transversions A-C (13%), G-T (13%), A-T (10%) and C-G (4%) (Fig. 1, A to C). These statistics were highly similar for the two metagenomes, with a comparable transition to transversion ratios of around 1.5. Yet, of all the SNVs identified, only 429,530 (54.83%) were common to the two metagenomes. Thus, local diatom populations can harbour a considerable density of genomic variability, the identity of which can vary dramatically between oceanic samples. Based on these results, we wondered how this extent of variability could emerge and be sustained over time.

**Fig. 1.**
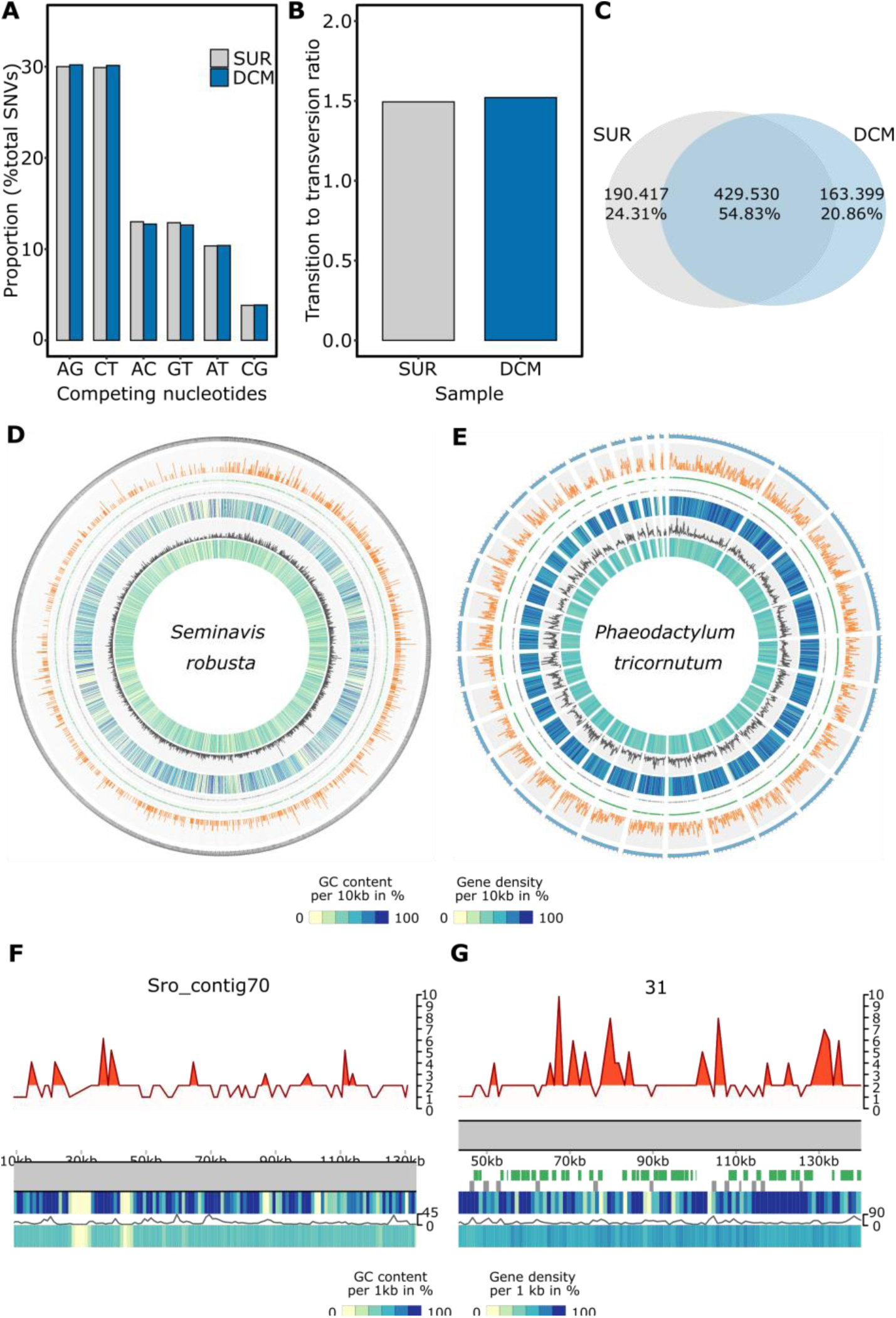
Intra-population variability of F. cylindrus in surface and DCM samples and genome-wide and locus-specific distribution of haplotypes in P. tricornutum and S. robusta. Overview of (**A**) the proportion of the competing nucleotides and (**B**) the transition to transversion ratio for surface and DCM samples. (**C**) Diagram representing the overlap between SNVs detected in the SUR and DCM metagenomes. SUR – surface, DCM - deep chlorophyll maximum. Genome-wide and locus-specific distribution of haplotypes. Summary of genome-wide detection of haplotypes per 1kb loci in (**D**) S. robusta contigs above 20 kb and (**E**) P. tricornutum chromosomes 1-33. From outside to inside: number of loci with more than two haplotypes* (orange), loci with two haplotypes (green), loci with a single haplotype (grey), gene density*, SNP density* (black), GC content*; * per 10 kb. (**F** to **G**) Example of a representative genomic region from (F) S. robusta and (G) P. tricornutum. The chromosome is represented by a grey rectangle. Above the chromosome: the number of observed haplotypes (orange line), loci with more than two haplotypes (orange area). Below chromosome from top to bottom: loci with two haplotypes (green), loci with a single haplotype (grey), gene density**, SNP density** (grey line), GC content**; ** per 1 kb.

Because the impact of sexual reproduction cannot be ruled out in natural populations, we examined haplotype diversity in clonal cultures from a single cell of two diatom model laboratory species: *P. tricornutum*, a species without observed sexual reproduction, and the obligate sexual heterothallic species *Seminavis robusta (8)*. Both were grown under conditions that preclude sexual reproduction. First, through sequence analysis, we measured the number of haplotypes per 1 kb in corrected PacBio and MinION sequencing data (Fig. S1). This analysis uncovered 1,405 loci with more than two haplotypes in *S. robusta* (125.57 Mb genome size) and 3,380 loci in *P. tricornutum* (27.4 Mb) (Fig 1, D to G, Data S2). Equivalent counting in haploid *Saccharomyces cerevisiae* (12 Mb) and diploid *Arabidopsis thaliana* (135 Mb) genomes yielded 3 and 83 loci with multiple haplotypes, respectively (Table S1 and Data S2). Further analysis of diatom loci with multiple haplotypes revealed that the GC content and SNP (single-nucleotide polymorphism, present in at least 20% of reads from the population) distribution were comparable to those in the rest of the genome (Fig. S2 and Table S2).

Genome-wide haplotype counting via long-read sequencing can suffer from increased sequencing noise, despite the correction, which could lead to an overcounting of haplotypes (Supplementary Text). We therefore validated the genome-wide data by observing selected loci from fresh, single-cell cultures (Fig. S3). For *S. robusta*, we profiled three loci identified in the genome-wide haplotype analysis in three independent cultures, four months after single diploid cell isolation. To overcome the potential problem of artefact generation during DNA amplification, we used emulsion PCR followed by Sanger sequencing of cloned PCR products. While the control mixture of two different alleles returned the two original haplotypes after PCR, we observed 2 to 6 haplotypes for the endogenous *S. robusta* loci (Fig. 2A, Table S3). In every case, two prominent haplotypes were supported by a higher number of reads, possibly representing the haplotypes present in the founder cell, whereas the additional haplotypes presumably appeared during the four months in culture.

**Fig. 2.**
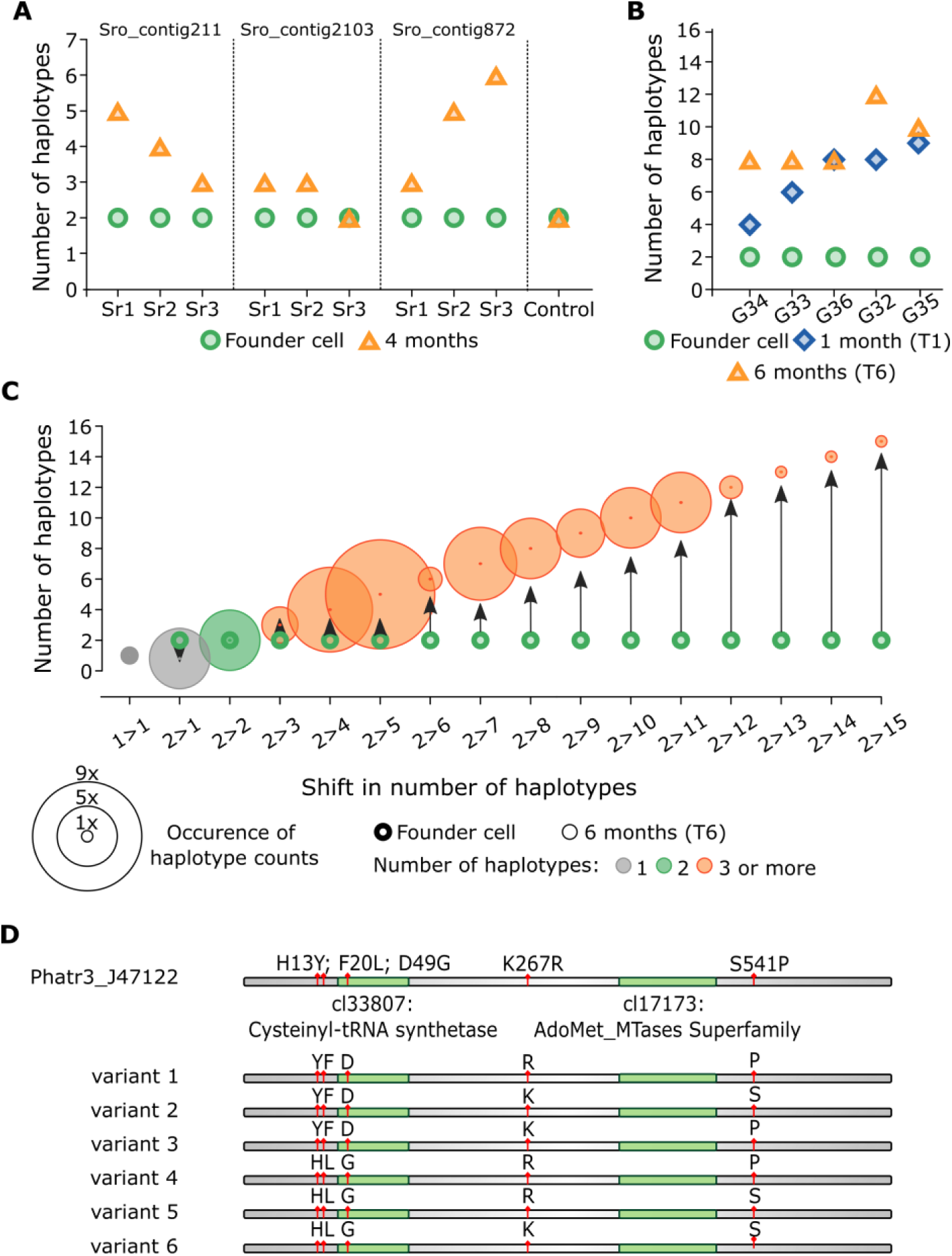
Accumulation of novel haplotypes in cultures freshly initiated from a single cells. (**A**) Detection of haplotypes on three loci in three S. robusta cultures four months after cultivation from a single cell. (**B** to **C**) Detection of the number of haplotypes in P. tricornutum at 1 month (T1) and 6 months (T6) after cultivation from a single cell. (B) Haplotypes at endogenous loci G32-G36 detected 1 and 6 months after single-cell isolation. (C) Change in the number of haplotypes at endogenous loci from founder cell (thick outline) to the number of haplotypes detected 6 months after single-cell isolation (thin outline). Categories of the shift in the number of haplotypes from founder cell to the number of haplotypes detected at T6 are on the x-axis, the number of haplotypes on the y-axis. The size of the circle corresponds to the number of cases in a given category, the color corresponds to the number of haplotypes; one haplotype (grey), two haplotypes (green), and three or more haplotypes (orange). (**D**) Schematic representation of predicted proteins variants in Phatr3_J47122 gene. Top line shows the position of amino acid variants on the protein indicated by red flags. Green regions depict conserved domains and lines below represent individual predicted variants.

Independently, haplotype diversity and the rate at which new haplotypes appeared were analyzed for 62 *P. tricornutum* 2-kb loci using emulsion PCR followed by PacBio amplicon sequencing. The heterozygosity of selected loci was profiled by SNP calling on the culture used for amplification, 1 month (T1) after single-cell isolation. Five loci (G32 to G36) were amplified 1 month (T1) and 6 months (T6) after single-cell isolation, whereas the remaining 57 loci were amplified at T6 only. The control reactions for random errors and artificial haplotype detection yielded the expected one and two haplotypes respectively, demonstrating that the emulsion PCR, PacBio library preparation, and sequencing did not generate artefacts (*9*) (Fig. S4A, Table S4). The number of recovered haplotypes varied between 1 and 15 (Fig. 2C, Fig. S4B and Table S5), with 6 loci displaying a single haplotype, 5 loci with two haplotypes, and 51 loci displaying at least three haplotypes. For four out of five loci amplified at both T1 and T6, an increase in the number of haplotypes was observed in the T6 sample (Fig. 2B, Fig. S4A) despite deeper sequencing coverage of the T1 samples, suggesting that haplotypes accumulate over time (Table S6). We analyzed the impact of haplotype variability on protein sequence in 20 genes fully covered by amplicon sequencing and found six for which the different haplotypes resulted in more than two putative protein variants, with up to six variants in the diatom-specific gene *Phatr3_J47122* (Fig. 2D, Fig. S5, Table S7).

Although only the locus G22 was identified as homozygous by SNP calling in T1, five additional loci with single haplotypes were found in T6 sequencing (Fig. S4A). These loci were identified as being heterozygous by SNP calling in T1, suggesting a loss of heterozygosity (LOH), which is generally associated with chromosomal rearrangements, gene conversion, and mitotic crossing-over (*10*). Moreover, the novel haplotypes that accumulated in both *S. robusta* and *P. tricornutum* cultures were recombinants lacking *de novo* mutations. Such new combinations are typically generated during sexual reproduction through meiotic recombination of homologous chromosomes (*11*). In contrast, interhomolog recombination is strictly suppressed in vegetative cells as it can result in harmful LOH events, chromosome rearrangements, and the onset of cancer in multicellular organisms (*12*). As sexual reproduction was excluded in our cultures, we tested whether haplotype diversity in clonal diatom populations might be the result of chromosomal rearrangements and interhomolog recombination in vegetative cells.

We sought to detect LOH and copy number variation (CNV) events in *P. tricornutum* under controlled conditions over a defined number of cell divisions. Three independent mother cultures (MC1 - MC3) were initiated from a single cell isolate and cultivated under conditions allowing approximately a single cell division per day (Fig. 3A, Fig. S6). After 30 days (T1), three single cells were again isolated from each mother culture to obtain nine daughter cultures (DC1.1 - DC3.3) that were harvested 30 days later (T2). At both T1 and T2, part of the mother cultures was also harvested. Following genome resequencing and SNP calling of all cultures, a pairwise comparison between the individual daughter cultures and their respective mother cultures was performed to identify novel CNVs and tracts of at least three consecutive SNPs that were lost in the daughter culture.

**Fig. 3.**
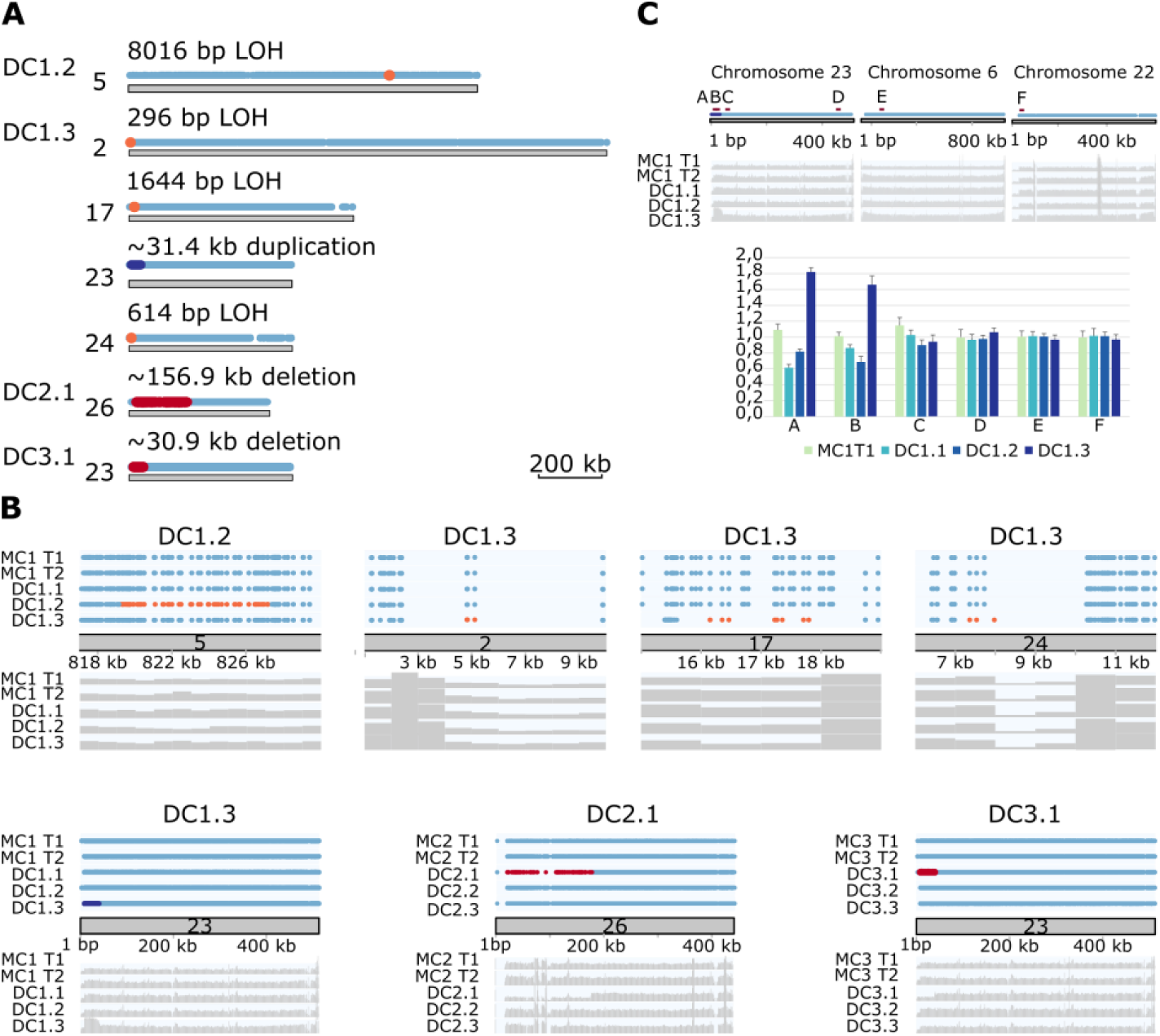
Genome-wide detection of LOH in P. tricornutum mother and daughter cultures after 30 days. (**A**) Position of copy-neutral LOHs (orange), duplication (dark blue) and deletions (red) in individual daughter cultures. Heterozygous regions are in light blue. Blank space – no SNPs. Numbers at the left refer to the chromosome number. (**B**) Zoomed-in regions with detected LOH, duplication or deletion events in the respective mother and daughter cultures. Blue dots – heterozygous SNPs, orange dots – homozygous SNPs in copy-neutral LOHs, red dots - SNPs in deletions, grey area – sequence coverage. (**C**) Confirmation of duplication on chromosome 23 in culture DC1.3 by qPCR. Position of target loci (red boxes) is shown in the upper part. Bar chart depicts the fold change in comparison to MC1T1 sample on control loci D, E and F. Blue dots – heterozygous SNPs, dark blue - SNPs in duplication, grey area – sequence coverage.

Events were found in four out of nine daughter cells. One copy-neutral 8016 bp LOH, where one allele of the locus was replaced by the other allele, was observed in DC1.2, three copy-neutral LOH events (296 bp, 614 bp and 1644 bp in length) and a 31.4 kb duplication covering 14 genes were observed in DC1.3 culture, and one 30.9 kb and one 156.9 kb deletion were detected in cultures DC3.1 and DC2.1, respectively, and confirmed by Sanger re-sequencing or qPCR (Fig. 3, Fig. S6A and B, Table S8 and S9). Besides the LOH events that were unique to a respective daughter culture, we identified several regions with reduced SNP density common to all cultures. SNP density was ten times lower than the genome average over almost the entire chromosome 19, and seventeen and thirty-eight times lower at the extremities of chromosomes 27 and 28, respectively, in comparison with the rest of the chromosome (Fig. S6C). These regions were not found to be SNP poor when sequencing the same *P. tricornutum* strain from other laboratories (*13, 14*), therefore we hypothesize that these regions represent early LOH events that occurred in one of the founder cells of our Pt1 subpopulation.

Profiling of the functional effect of SNPs in LOH regions revealed 18 SNPs with possible high impacts on gene function, including 3 SNPs that caused a loss of function by introducing a premature stop codon in the respective gene (Table S10).

While the mechanism behind the observed deletions and duplication remains difficult to interpret, such events can happen through non-allelic homologous recombination or unequal mitotic crossing over between both sister chromatids or homologous chromosomes (*15, 16*). In contrast, copy-neutral LOH events require an exchange of genetic information between homologous chromosomes. To estimate the rate of interhomolog recombination in *P. tricornutum*, we established a tractable endogenous readout system for copy-neutral LOH detection, based on three strains containing two different mutant alleles of the *PtUMPS* gene, generated through gene editing (*17, 18*). In strain *ptumps-1bp*, the point mutations in the two alleles occur at a position only 1 bp apart, in strain *ptumps-320bp* they are separated by 320 bp and in strain *ptumps-1368bp* by 1368 bp (Fig. 4A). As the PtUMPS protein is required for uracil biosynthesis, cells with a wild-type (WT) allele can synthesize uracil, but also convert 5-fluoroorotic (5-FOA) acid into the toxic 5-fluorouracil (5-FU), resulting in cell death. In contrast, mutant cells are resistant to 5-FOA but are uracil auxotrophs. The *ptumps-/-* strains were cultivated under non-selective conditions for 14 days (with uracil and without 5-FOA) to permit potential recombination at the *PtUMPS* locus (Fig. S7). Subsequently, 5×10^7^ cells from the culture were plated on medium without uracil to select cells that underwent recombination at the *PtUMPS* locus and restored the WT allele. We recovered no colonies in strain *ptumps-1bp*, 12 colonies in strain *ptumps-320bp* and 83 colonies in strain *ptumps-1368bp* (Fig. 4B, Fig. S8, Table S11). The lack of colonies in the *ptumps-1bp* strain confirmed that the WT allele was not restored by a random mutation. Moreover, sequencing of *PtUMPS* alleles from ten *ptumps-1368bp* colonies and five *ptumps-320bp* colonies confirmed the restoration of the WT allele through copy-neutral LOH events (Fig. 4C, Fig. S7).

**Fig. 4.**
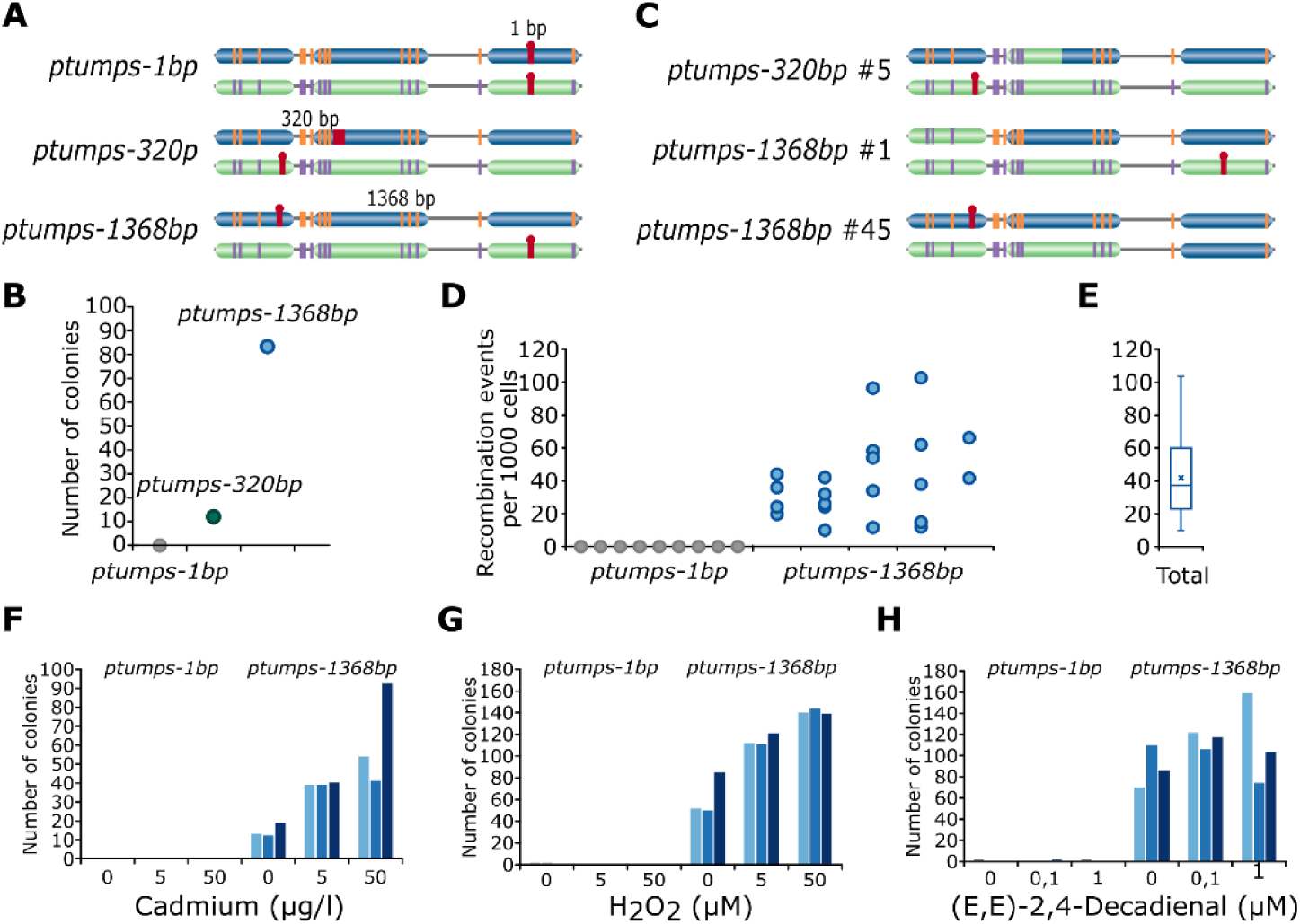
Detection of LOH events at the P. tricornutum PtUMPS locus. (**A**) Schematic of alleles in the PtUMPS strains. Homologous chromosomes are depicted in blue and green, loss-of-function mutations are in red. (**B** to **C**) Recombination in PtUMPS mutant strains during 14 days of cultivation under non-selective conditions. (**B**) The number of recovered uracil prototrophic colonies per strain. (**C**) Examples of sequenced recombinant alleles in one ptumps-320bp and two ptumps-1368bp colonies. (**D** to **E**) Estimation of the interhomolog recombination frequency. (D) Recombination rate per thousand cell divisions. Each dot represents one replica. (E) The box plot shows from bottom to top: minimum value, first quartile, median (line), mean (cross), third quartile, and maximum value. (**F** to **G**) Recombination events in response to stress-induced by (F) cadmium; (G) H_2_O_2_; (H) 2,4-Decadienal. ptumps-1368bp replicas are depicted in different shades of blue.

Next, the *PtUMPS* system was used to obtain an estimate of the interhomolog recombination frequency. A total of 2×10^7^ cells from 5-FOA- and uracil-supplemented medium were directly plated onto medium without uracil to select only those cells that were in the process of interhomolog recombination during a single round of cell division. The frequency of interhomolog recombination was 4.2 per 100 cell divisions (Fig. 4D and E, Fig. S7 and S9 to S11, Table S12), approximately ten times higher than the rate reported for *S. cerevisiae* after recalculation per cell division (*19, 20*).

To test whether the rate of interhomolog recombination can be influenced by environmental conditions, we employed the *PtUMPS* readout system to test the effect of three physiologically relevant stresses: hydrogen peroxide (H_2_O_2_), which is produced by various phytoplankton groups and can act as a signaling molecule as well as cause oxidative damage (*21*); the trace metal cadmium (*22*), which contaminates aquatic environments; and a polyunsaturated aldehyde (E,E)-2,4-Decadienal which is involved in diatom intercellular signaling, stress surveillance, and defense against grazers but which can trigger lethality at high concentrations (*23, 24*). For each mock and stress treatment, 25×10^6^ cells per replica were transferred from 5-FOA- and uracil-supplemented medium (preventing recombination at the *PtUMPS* locus) to medium containing uracil for 24 h, thus allowing a maximum of one cell division. Next, cells were plated on selective medium without uracil to recover cells that restored the WT *PtUMPS* allele through interhomolog recombination. Whereas (E,E)-2,4-Decadienal treatment did not influence the rate of interhomolog recombination, we found a positive, concentration-dependent effect of H_2_O_2_ and cadmium on the number of colonies recovered (Fig. 4F to G, Fig. S12 to S17, Tables S13 to S15). These data illustrate that environmental stresses increase the frequency of recombination between homologous chromosomes.

Our results demonstrate an unusually high occurrence of chromosomal rearrangements and mitotic interhomolog recombination in *P. tricornutum*, resulting in the rapid accumulation of new haplotypes and fixation of polymorphisms in a homozygous state. The capability to rapidly fix novel SNVs in a population through LOH could explain the differences observed in the metagenomes of *F. cylindrus* subpopulations from surface and DCM samples from the same station. Preceding studies highlighted extensive allelic diversity in the pennate polar diatom *F. cylindrus (3)* and the presence of large low-SNP-density regions, indicative of LOH events, in the otherwise highly heterozygous genome of the centric diatom *T. pseudonana (6)* as well as another *P. tricornutum* Pt1 subpopulation (*25*). This suggests that the phenomenon of frequent chromosomal rearrangement and interhomolog recombination, previously detected in pathogenic oomycetes *(26)* and yeasts (*20, 27*), may be widespread across diatoms.

The life cycle of many diatom species is characterized by long periods of clonal reproduction, thus the creation of novelty and recombinant genotypes is limited. Genomic rearrangements such as duplications represent a major driving force in the gain of new biological functions (*28*), whereas interhomolog mitotic recombination generates novel allelic combinations within clonal lineages. While the gain of new functions through gene duplication requires the emergence of novel mutations, the variability of haplotypes at the genomic level can be immediately translated into differences in the proteome and potentially result in physiological divergence. Indeed, according to Wright’s classical theory of dominance (*29, 30*), a considerable amount of phenotypic remains hidden as recessive variation, locked away behind the dominant alleles at the same loci. The increase in the frequency of interhomolog mitotic recombination rate under exposure to stress can convert this non-additive genetic variation into a variation that is instantly available to natural selection, which may further boost the divergence of the clonal population during a critical period and enhance the adaptive potential of clonally propagated diatom populations to respond to environmental changes. Furthermore, fixation of mutations in sexual reproduction genes or allopatric populations, possibly combined with adaptation to local conditions (*31, 32*), could contribute to lineage diversification and ultimately to the speciation. Thus, we hypothesize that evidence presented here, both from a natural diatom population and from clonally propagated species in the lab, for the unusually high frequencies of chromosomal rearrangements and interhomolog mitotic recombination in diatoms may underlie the ecological success of diatoms by creating a high level of genetic variability within clonal lineages upon which natural selection can act.

## Acknowledgments

We thank Dr. Nicole Poulsen for providing GK-359 and GK-333 plasmids and Dr. Annick Bleys and Dr. James Matthew Watson for proofreading the manuscript.

## Funding

This research was supported by BOF project GOA01G01715 to L.D.V., K.V. and W.V.; Erwin Schrödinger fellowship from Austrian Science Fund (project J3692-B22) to P.B.; Gordon and Betty Moore Foundation grant (GBMF 4966), a Région Midi-Pyrénées grant (15058490 financial support for Accueil d’Equipes d’Excellence), an ANR JCJC grant (ANR-16-CE05-0006-01), and the 3BCAR Carnot Institute funding to D.J. and F.D.; C.N. and C.B. were supported by European Research Council (ERC) under the European Union’s Horizon 2020 research and innovation programme through the project DIATOMIC (grant agreement No. 835067) to C.B.

## Author contributions

P.B. and L.D.V. conceptualized the study. P.B. initiated, designed and performed all experiments and bioinformatic analysis in the manuscript. M.S. performed sequencing of uracil prototrophic colonies. D.J. generated *ptumps* mutant strains. C.N and T.D. performed the *F. cylindrus* metagenome assembly and analysis, I.V. helped with diatom culture maintenance. C.M.O-C. and E.M. provided bioinformatic datasets. P.B. wrote the manuscript, generated all figures and data visualizations. L.D.V., K.V., F.D., C.B., W.V., K.S. supervised the research. P.B., L.D.V, K.V., W.V., F.D, K.S., C.B., C.V.O., T.M. reviewed and edited the manuscript.

## Competing interests

The authors declare no competing interests.

## Data and materials availability

Raw sequencing data were deposited to the Sequence Read Archive (SRA) under BioProject accessions PRJNA658511 and PRJNA658224. SRA accession numbers for individual samples are listed in Supplementary Data Table 3. Processed datasets were uploaded to zenodo: Aligned and processed long-read sequencing datasets *S. robusta* PacBio, *P. tricornutum* MinION reads and SNP selected for haplotype counting for both species are available at https://doi.org/10.5281/zenodo.4005721. Aligned PacBio amplicon sequencing reads, reference file and selected biallelic SNPs used in haplotype counting are available at https://doi.org/10.5281/zenodo.4005643. Processed datasets from LOH detection in mother and daughter cultures including ILLUMINA reads aligned to the reference P. tricornutum genome used for SNP calling, SNP calls for individual samples and jointly called SNPs on all samples are available at https://doi.org/10.5281/zenodo.4006016. The haplotype coding script to count haplotypes in long-read sequencing datasets is available on zenodo, at https://doi.org/10.5281/zenodo.4001752. A version of the haplotype counting script that outputs the combination of bases at selected SNP sites is available on zenodo, at https://doi.org/10.5281/zenodo.4173002. All other data are available from the authors upon request.

## Supplementary Materials (separate file)

Materials and Methods

Supplementary Text

Figs. S1 to S17

Tables S1 to S16

References (31-64)

Data S1 (separate file)

Data S2 (separate file)

Data S3 (separate file)

## Notes

### Competing Interest Statement

The authors have declared no competing interest.

## References and Notes

1. S. Malviya et al., Insights into global diatom distribution and diversity in the world’s ocean. P Natl Acad Sci USA 113, E1516–E1525 (2016).

2. A. Godhe, T. Rynearson, The role of intraspecific variation in the ecological and evolutionary success of diatoms in changing environments. Philos T R Soc B 372, (2017).

3. T. Mock et al., Evolutionary genomics of the cold-adapted diatom Fragilariopsis cylindrus. Nature 541, 536–540 (2017).

4. M. Krasovec, S. Sanchez-Brosseau, G. Piganeau, First Estimation of the Spontaneous Mutation Rate in Diatoms. Genome Biol Evol 11, 1829–1837 (2019).

5. V. A. Chepurnov et al., In search of new tractable diatoms for experimental biology. Bioessays 30, 692–702 (2008).

6. J. A. Koester et al., Sexual ancestors generated an obligate asexual and globally dispersed clone within the model diatom species Thalassiosira pseudonana. Sci Rep-Uk 8, (2018).

7. W. M. Lewis, The Diatom Sex Clock and Its Evolutionary Significance. American Naturalist 123, 73–80 (1984).

8. V. A. Chepurnov, D. G. Mann, W. Vyverman, K. Sabbe, D. B. Danielidis, Sexual reproduction, mating system, and protoplast dynamics of Seminavis (Bacillariophyceae). J Phycol 38, 1004–1019 (2002).

9. Materials and methods are available as supplementary materials at the Science website.

10. M. Hiraoka, K. Watanabe, K. Umezu, H. Maki, Spontaneous loss of heterozygosity in diploid Saccharomyces cerevisiae cells. Genetics 156, 1531–1548 (2000).

11. N. Hunter, Meiotic Recombination: The Essence of Heredity. Cold Spring Harb Perspect Biol 7, (2015).

12. A. J. R. Bishop, R. H. Schiestl, Homologous recombination as a mechanism of carcinogenesis. Bba-Rev Cancer 1471, M109–M121 (2001).

13. A. Rastogi et al., A genomics approach reveals the global genetic polymorphism, structure, and functional diversity of ten accessions of the marine model diatom Phaeodactylum tricornutum. Isme J 14, 347–363 (2020).

14. National Library of Medicine (US), in Bethesda (MD): National Center for Biotechnology Information. (2018).

15. I. Alves, A. A. Houle, J. G. Hussin, P. Awadalla, The impact of recombination on human mutation load and disease. Philos Trans R Soc Lond B Biol Sci 372, (2017).

16. L. S. Symington, R. Rothstein, M. Lisby, Mechanisms and Regulation of Mitotic Recombination in Saccharomyces cerevisiae. Genetics 198, 795–835 (2014).

17. T. Sakaguchi, K. Nakajima, Y. Matsuda, Identification of the UMP Synthase Gene by Establishment of Uracil Auxotrophic Mutants and the Phenotypic Complementation System in the Marine Diatom Phaeodactylum tricornutum. Plant Physiol 156, 78–89 (2011).

18. M. Serif et al., One-step generation of multiple gene knock-outs in the diatom Phaeodactylum tricornutum by DNA-free genome editing. Nat Commun 9, (2018).

19. M. A. Barbera, T. D. Petes, Selection and analysis of spontaneous reciprocal mitotic cross-overs in Saccharomyces cerevisiae. P Natl Acad Sci USA 103, 12819–12824 (2006).

20. Y. Sui et al., Genome-wide mapping of spontaneous genetic alterations in diploid yeast cells. Proc Natl Acad Sci U S A, (2020).

21. B. D’Autreaux, M. B. Toledano, ROS as signalling molecules: mechanisms that generate specificity in ROS homeostasis. Nat Rev Mol Cell Bio 8, 813–824 (2007).

22. T. Brembu, M. Jorstad, P. Winge, K. C. Valle, A. M. Bones, Genome-Wide Profiling of Responses to Cadmium in the Diatom Phaeodactylum tricornutum. Environ Sci Technol 45, 7640–7647 (2011).

23. A. Ianora et al., Aldehyde suppression of copepod recruitment in blooms of a ubiquitous planktonic diatom. Nature 429, 403–407 (2004).

24. A. Vardi et al., A stress surveillance system based on calcium and nitric oxide in marine diatoms. Plos Biol 4, 411–419 (2006).

25. M. T. Russo, R. A. Cigliano, W. Sanseverino, M. I. Ferrante, Assessment of genomic changes in a CRISPR/Cas9 Phaeodactylum tricornutum mutant through whole genome resequencing. Peerj 6, (2018).

26. A. L. Dale et al., Mitotic Recombination and Rapid Genome Evolution in the Invasive Forest Pathogen Phytophthora ramorum. Mbio 10, (2019).

27. A. Gusa, S. Jinks-Robertson, Mitotic Recombination and Adaptive Genomic Changes in Human Pathogenic Fungi. Genes (Basel) 10, (2019).

28. J. Z. Zhang, Evolution by gene duplication: an update. Trends Ecol Evol 18, 292–298 (2003).

29. S. Wright, Fisher’s Theory of Dominance. The American Naturalist 63, 274–279 (1929).

30. S. Wright, Physiological and Evolutionary Theories of Dominance. The American Naturalist 68, 24–53 (1934).

31. E. Pinseel et al., Global radiation in a rare biosphere soil diatom. Nat Commun 11, (2020).

32. T. Y. James et al., Adaptation by Loss of Heterozygosity in Saccharomyces cerevisiae Clones Under Divergent Selection. Genetics 213, 665–683 (2019).

