## Supplementary Data for "Mitotic interhomolog recombination drives genomic diversity in diatoms"

**This PDF file includes:**

Materials and Methods
Supplementary Text
Figs. S1 to S17
Tables S1 to S16
References

**Other Supplementary Materials for this manuscript include the following:**

Data S1 (Excel)
Data S2 (Excel)
Data S3 (Excel)

### Materials and Methods

Estimation of intra-specific variability in *Fragilariopsis cylindrus* metagenomes

*Tara* Oceans metagenomic reads from 0.8-5 µm, 5-20 µm, 20-180 µm, 180-2000 µm and 0.8-2000 µm size fractions were mapped against the FASTA file of the *Fragilariopsis cylindrus* CCMP 1102 genome (*31*) (available at http://genome.jgi-psf.org/Fracy1/Fracy1.home.html) using Bowtie2 v2.3.4.3 (*32*) with a 95% identity filter. Two depths (surface and deep chlorophyll maximum, DCM) from Station 86, located in the Southern Ocean (near the Antarctic peninsula, 64°30'88" S, 53°05'75" W), displayed vertical coverage superior or equal to 10X for three size fractions (0.8-5, 5-20 and 0.8-2000) and were then selected for further analysis. Using SAMtools v1.10 (*33*), the resulting SAM files were converted into BAM files and for each sample the BAM files of the three different size fractions were merged to increase the coverage (final mean coverage 51.75X and 58.46X for surface and DCM). Downstream analyses were performed with the anvi'o platform (*34*) to generate profile databases based on the BAM files that were combined into a merged profile database. Genes were imported into anvi'o at the level of individual exons. Then, the program "anvi-summarize" was used with the "init-gene-coverages" flag to characterize the mean coverage of each gene in the surface and DCM samples. A subset of *F. cylindrus* genes was defined as the core genes (n = 24,326) if they occurred in the two samples and their mean coverage in each sample remained within a factor 3 of the mean coverage of all 27,137 genes in the same metagenome. This step prevented the analysis of genes lacking signal or recruiting reads from other species (*35*). Finally, the intra-population variability of *F. cylindrus* was analysed across the core genes and in the two samples using the programme "anvi-gen-variability-profile", which provided tables reporting single nucleotide variants (hereafter referred to as SNVs) and their nucleotide frequencies in the recruited reads. We defined SNVs as positions displaying at least 10% variation from the consensus nucleotide and with a mean vertical coverage ≥ 20X in the two samples. The variability tables were imported into R v4.0.1 to compute the number of variable positions and SNV density (i.e. the number of positions with SNVs for each exon in the core genes divided by the corresponding exon length) for each exon. Gene-level mean coverage, number of variable positions and SNV density were computed using the information from the individual exons.

Diatoms datasets and strains

Datasets and strains used in this study are summarized in Data S3. The S. robusta D6 reference strain (accession number DCG 0498) is available from the BCCM/DCG diatom culture collection at Ghent University (http://bccm.belspo.be/about-us/bccm-dcg). Publicly available genomes of *S. robusta* strain D6 (*36*) https://www.ebi.ac.uk/ena/browser/view/CAICTM010000000 and *P. tricornutum* Pt1 8.6 (CCMP2561) strain (*37*) https://www.ebi.ac.uk/ena/browser/view/GCA_000150955.2 and next-generation sequencing datasets were used for our analysis.

Genome-wide haplotype counting in *S. robusta* and *P. tricornutum* next-generation sequencing data

For both *S. robusta* and *P. tricornutum* genome-wide haplotype counting, a reliable single nucleotide polymorphism (hereafter referred to as SNP; in contrast to SNVs found in metagenomes from natural populations, SNP had be supported by at least 20% of reads in sample from laboratory single strain) set was first identified in ILLUMINA short-read sequencing datasets and then used for counting of the number of haplotypes in the PacBio RS II and MinION long reads. The ILLUMINA and PacBio data of *S. robusta* from https://www.ebi.ac.uk/ena/browser/view/PRJEB36614 and ILLUMINA and Minion data of *P. tricornutum* from https://www.ebi.ac.uk/ena/browser/view/PRJNA487263. Because in long-read sequencing the error rate for indels is higher than for SNPs, indels were ignored in our analysis.

*SNP calling*: SNP calling on ILLUMINA short-read sequencing was done using GATK HaplotypeCaller 3.7.0 (*38*). In short, adapters and reads with the quality score below 20 were removed from ILLUMINA reads using BBduk2 (*39*) with minlen=35 qtrim=rl trimq=20 hdist=1 tbo tpe options and custom adapter reference file. Next, the respective reads were aligned to *S. robusta* v1 assembly CAICTM010000001-CAICTM010004752 (European Nucleotide Archive) or *P. tricornutum* v2 assembly2 GCA_000150955.2 (European Nucleotide Archive) using Burrows-Wheeler Alignment Tool (BWA) (*40*) algorithm BWA-MEM with -M option. Unmapped and multi-mapped reads were removed using SAMtools (*33*) view with -h -F 4 -q 1 options. Aligned reads were then sorted using picard-tools 1.8.0 (*41*) SortSam and duplicate reads were marked with MarkDuplicates and indexed with BuildBamIndex. Read base quality scores were adjusted by two round of recalibration. Here, SNPs and indels were called by GATK HaplotypeCaller (*42*) and filtered with a set of hard filters using SelectVariants; QD < 2.0, FS > 60.0, MQ < 40.0, MQRankSum < -12.5, ReadPosRankSum < -8.0 for SNPs and QD < 2.0, FS > 200.0, ReadPosRankSum < -20.0 for indels. Recalibration table was generated with BaseRecalibrator and recalibrated reads were printed with PrintReads. After a second round of recalibration, germline SNPs were called using HaplotypeCaller with --genotyping_mode DISCOVERY. Next, reliable biallelic SNPs were selected using SelectVariants with --restrictAllelesTo BIALLELIC -selectType SNP and QD < 2.0, QUAL < 30.0, SOR > 3.0, FS > 60.2, MQ < 40.0, MQRankSum < -12.5, ReadPosRankSum < -8.0, AF > 0.2 and DP < 10 options.

Repeat regions and low complexity DNA sequences in *S. robusta* and *P. tricornutum* were identified using RepeatModeler 1.0.9 (*43*) and masked using RepeatMasker 4.0.5 (*44*), and SNPs in these regions were removed from the dataset using BEDtools (*45*) subtract algorithm. Finally, selected fields (CHROM, POS, REF, ALT) from the SNP dataset were extracted from the vcf file to a table and split into independent files by contig/chromosome using awk.

*S. robusta PacBio reads processing:* Circular Consensus Sequences (CCS) were obtained with smrtanalysis 2.3.0 (PacBio) with minFullPasses 0 option. CCS reads were then self-corrected using canu 1.4 (*46*) with canu_correct genomeSize=136.0m errorRate=0.035 -pacbio-raw options and trimmed with canu_trim genomeSize=136.0m errorRate=0.035 -pacbio-corrected options. Corrected reads were mapped to the reference genome using BLASR (*47*) with -sam -clipping soft options. The CIGAR string was corrected with samfixcigar, soft-clipped bases were removed with biostar84452 from jvarkit (*48*) and uniquely mapped reads with mapping quality >20 were selected using SAMtools (*33*). Coverage was estimated using GATK 3.7.0 DepthOfCoverage. SAMtools view and awk were used to split the PacBio reads to separate files per contig.

*P. tricornutum MinION reads processing*: MinION reads were self-corrected using canu 1.4 (*46*) with canu_correct genomeSize=30m errorRate=0.144 -nanopore-raw options. Reads were aligned to the genome using GraphMap (*49*) with default settings and uniquely mapped reads were selected using SAMtools view. The CIGAR string was corrected with samfixcigar, soft-clipped bases were removed with biostar84452 from jvarkit. Coverage was estimated using GATK 3.7.0 DepthOfCoverage. SAMtools view and awk were used to split the PacBio reads to separate files per contig.

*Haplotype counting:* The haplotype counting was done with a custom script based on bash, awk and Sam2Tsv from jvarkit (https://doi.org/10.5281/zenodo.4001752). In short, a record for every base in each processed PacBio/MinION read with the position, reference and the actual base was obtained using Sam2Tsv from jvarkit (*48*). Next, only positions of SNPs selected in ILLUMINA reads were retained. The record was divided into fixed windows of max 1 kb from the first SNP and haplotypes for selected sites were written for each read separately. Reads containing an indel or another base than the reference or the alternative base at the selected SNP position or not covering the 1 kb region were removed and the number of haplotypes and number of supporting reads for each haplotype was counted using awk. Loci with multiple haplotypes were selected from the record of the number of haplotypes per 1 kb with the number of supporting reads with following conditions: at least three haplotypes had to be supported each by at least 2 reads, the locus had to be at least 100 bp long and the coverage had to be below 100x in order to remove repeat regions that were not masked. Visualization of haplotype counting data was done using Circos (*50*) and karyoploteR (*51*).

*Haplotype counting in control genomes*: As a control for haplotype counting, the genome assembly, ILLUMINA and PacBio sequencing data publicly available for haploid yeast *Saccharomyces cerevisiae* GLBRCY22-3 (*52*) grown from single colony https://www.ncbi.nlm.nih.gov/bioproject/PRJNA279877 and for several diploid *Arabidopsis thaliana* Ler plants (*53, 54*) from https://www.ncbi.nlm.nih.gov/bioproject/PRJNA311266 and https://www.ncbi.nlm.nih.gov/bioproject/237120 were used. The SNP calling, PacBio data processing and haplotype counting were done as described above and are summarized in Table S1 and Data S2.

Resequencing of *S. robusta* and *P. tricornutum* loci with multiple haplotypes

*Diatom cultivation conditions:* Both *S. robusta* strain D6 and *P. tricornutum* strain Pt1 subculture MC2 were cultivated in 1x TMB medium consisting of 34.5 g/l of Tropic Marin Bio-Actif sea salt (Tropic Marin, Germany) and 0.08g/l sodium bicarbonate (Sigma-Aldrich) supplemented with 1x Guillard's (F/2) Marine Water Enrichment Solution (Sigma-Aldrich), 100 µg/ml ampicillin, 50 µg/ml gentamycin and 100 µg/ml streptomycin in 12 h/12 h light/dark cycle. *P. tricornutum* cultures were cultivated at 20°C, under photosynthetic LED light with an intensity of 160 µmol photons $m^{-2}$ $s^{-1}$ and with 100 rpm shaking. *S. robusta* cultures were cultivated at 18°C with approximately 85 µmol photons $m^{-2}$ $s^{-1}$ from cool-white fluorescent lights.

*DNA extraction:* DNA for deep sequencing was harvested and isolated by the CTAB DNA extraction method. Cells from approximately 500 ml of exponentially growing *S. robusta* and *P. tricornutum* cultures were harvested by centrifugation at 1216 x g for 5 min. The supernatant was discarded and cell pellet was resuspended in 400 µl of CTAB buffer (1% (w/v) CTAB, 100 mM Tris-HCl pH 7.5, 10 mM EDTA pH 8, 700 mM NaCl and freshly added 4 µg RNase A). *S. robusta* cells were disrupted by agitation with glass/zirconium beads (0.1-mm diameter; Biospec) on a bead mill (Retsch) for three times 1 min at frequency 20 Hz. Samples were incubated for 30 min at 60°C and afterwards let to cool down on ice for 15 min. Next, 250 µl of chloroform:isoamylalcohol 24:1 was added and the samples were mixed manually for 1 min. Phases were separated by centrifugation at 20 000 x g for 10 min. The upper aqueous phase was transferred to a new tube and DNA was precipitated by addition of an equal volume of isopropanol followed by centrifugation for 15 min at 20 000 x g. The DNA pellet was washed with 70% ethanol, air-dried and resuspended in 50 µl of 10 mM Tris-HCl pH 8.5.

*Emulsion PCR:* Loci for re-sequencing of haplotypes were selected from the list of loci with multiple haplotypes obtained through genome-wide haplotype detection. Three loci were selected in *S. robusta* for Sanger sequencing verification (Table S3) and 62 loci for PacBio amplicon sequencing verification in case of *P. tricornutum* (Table S5). Primers for amplification were designed manually (Data S3). To avoid PCR recombination artefacts (*55, 56*), selected loci were amplified by emulsion PCR using the MICELLULA DNA Emulsion & Purification Kit (roboklon, Germany) according to the manufacturer's instructions. The DNA concentration was measured on NanoDrop (ThermoFisher Scientific) and the number of DNA template copies per µg of DNA was calculated according to the genome size of the respective diatom. Maximum of $10^7$ of DNA molecules were used per single emulsion PCR reaction. The PCR reaction mix consisted of 1x OptiTaq PCR buffer B, 200 µM dNTP mix, 2 µM of each forward and reverse primer, DNA template with $10^6$-$10^7$ molecules, 1 mg/ml acetylated BSA and 2.5U Opti Taq DNA polymerase in 50 µl of total volume. Emulsion mix was prepared separately by mixing 220 µl of emulsion component 1, 20 µl of emulsion component 2 and 60 µl of emulsion component 3 per PCR reaction. The 50 µl PCR reaction was mixed with 300 µl of emulsion mix and emulsion was created by continuous vortexing at 1400 rpm at 4°C for 5 min. Each emulsion PCR reaction was split into three PCR tubes and run with following parameters: 94°C initial denaturation for 2 min, 26 cycles of 94°C denaturation for 15 s, 56°C annealing for 30 s and 72°C extension with 1 kb/min relative to the amplified fragment length, followed by a final extension at 72 °C for 10 min. The emulsion was broken by the addition of 1 ml of isobutanol and vortexing. Next, 400 µl of Orange-DX solution was added and reactions were gently mixed and centrifuged for 2 min at 20 000 x g. The organic phase was removed and the aqueous phase was transferred to a Micellula spin column activated by 40 µl of DX buffer. Columns were centrifuged at 11 000 x g for 1 min, washed first with 500 µl of Wash-DX1 buffer, and then with 650 µl of Wash-DX2 buffer and the leftovers of buffer were removed by an additional centrifugation for 2 min. PCR products were eluted in 50 µl of Elution-DX buffer (all components: roboklon).

Sanger sequencing of *S. robusta* amplicons

*S. robusta* emulsion PCR products were cloned into the pGEM-T vector (Promega) according to the manufacturer's instructions. In brief, the A overhangs were added by incubation of PCR product with 10 µM dNTP mix and 1U of Taq DNA polymerase (Invitrogen) at 72°C for 10 min. Then, 3.5 µl of PCR product was mixed with 5 µl of 2x ligase buffer, 0.75 µl of pGEM-T vector and 2.25 U of T4 DNA ligase (all Promega) and incubated for 12 h at 4°C. Ligation mixtures were transformed through electroporation into *E. coli* DH5alpha cells and transformants were selected on LB supplemented with 100 µg/ml ampicillin (Duchefa). Clones containing cloned PCR products were selected by Sanger sequencing with pGEM-5 and pGEM-6 primers (Data S3). Sequencing results were aligned with the reference using Clustal Omega (*57*), and haplotypes were manually assembled for each clone. Two alleles of Sro_contig556:54453-55487 (Data S3) were cloned into the pGEM-T vector and an equimolar mix of these two plasmids was used for emulsion PCR as control for artefact generation. To simulate conditions similar to emulsion PCR reactions on *S. robusta* genomic DNA, $10^6$-$10^7$ molecules of *S. robusta* genomic DNA were added and the control samples were amplified with pGEM-3 and pGEM-4 primers (Data S3). The PCR products were again cloned into the pGEM-T vector and sequenced with pGEM-5 and pGEM-6 primers.

<u>*P. tricornutum* cultures and PacBio amplicon sequencing and haplotype counting</u>

13 intergenic loci and 59 loci overlapping with coding regions (Table S5) were selected based on an SNP call on a T1 cell culture and the list of loci with more than two haplotypes for PacBio Sequel amplicon sequencing. All loci were amplified by emulsion PCR as described above with primers listed in Data S3. In the case of low amplification efficiency, the emulsion PCR was repeated. Purified PCR products were concentrated using Genomic DNA Clean & Concentrator (Zymo Research) according to the manufacturer's instructions. Amplifications of *CFP*, *GFP* and *YFP* genes were used for control reactions for random mistakes and control reactions for artificial haplotypes detection. The *CFP*, *GFP* and *YFP* sequences (Table S4 and Data S3) were amplified by emulsion PCR with primers binding to vector backbone (Data S3) from GK-333-CFP, GK-359 and GK-333-YFP plasmids respectively (GK-333 and GK-359 were a gift from Dr. Nicole Poulsen, Center for Molecular Bioengineering at TU Dresden). Control reactions for random errors consisted of separately amplified GFP and YFP and control reactions for PCR-mediated recombination consisted of mixed amplification of CFP+GFP and CFP+YFP (Table S4). Amplicons were pooled together into two samples. Sample 1 contained 63 *P. tricornutum* endogenous 63 amplicons from DNA harvested at T6 time point, YFP amplified separately and CFP+GFP amplified in one reaction. Sample 2 contained 5 *P. tricornutum* endogenous 5 amplicons from DNA harvested at T1 time point, GFP amplified separately and CFP+YFP amplified in one reaction. Samples were barcoded, mixed in 9:1 ratio and sequenced on 1 PacBio Sequel SMRT cell at Novogene (UK).

Circular Consensus Sequence (CCS) were obtained with SMRT Link 8.0.0 software (PacBio) with --min-length 500 --max-length 2400 --min-passes 4 --by-strand and mapped to the reference loci with BLASR(*47*) with default settings. The CIGAR string was corrected with samfixcigar, soft-clipped bases were removed with biostar84452 from jvarkit (*48*) and reads with mapping quality >20 were selected using SAMtools (*33*). Next, haplotype number per locus was counted as described above. Only haplotypes supported either by at least 1% of valid reads or at least two reads if read count was lower than 200 were selected (Table S5). The genes fully covered by PacBio amplicons were manually annotated to detect putative variants.

<u>LOH and CNV detection</u>

*Cultures started from a single cell*: To isolate single cells from *P. tricornutum* cultures, an aliquot from the respective culture was diluted to $10^6$, $10^9$ and $10^{12}$ in the growth medium. 200 µl of diluted culture were spread on a 120 x 120 mm Petri dish (Corning Gosselin) with solid medium prepared with 17,25 g/l of Tropic Marin Bio-Actif sea salt solid and 10 g/l of Plant Tissue Culture Agar (Neogen) supplemented with 1x Guillard's (F/2) Marine Water Enrichment Solution (Sigma-Aldrich), 100 µg/l ampicillin, 50 µg/l gentamycin and 100 µg/l streptomycin, and the plates were incubated at room temperature with a 12-h/12-h light/dark cycle. Plates were checked for the presence of colonies after 14 days and single colonies were transferred from the plate with the highest dilution factor that contained colonies into liquid TMB medium using a pipette tip. This procedure was used to isolate and start three colonies from a single cell from the Pt1 culture to obtain mother cultures MC1, MC2 and MC3. Thirty days after mother culture isolation (T1 time point), daughter cells DC1.1-DC3.3 were isolated from the respective mother cultures (Fig. S6). Cultures for deep sequencing were harvested twice; on time point T1 part of the mother cultures was harvested at time point T1, and 30 days later at time point T2 all cultures in the experiment were harvested.

*Illumina sequencing, SNP calling, LOH and CNV detection:* DNA for deep sequencing was harvested and isolated by the CTAB DNA extraction method as described above. Paired-end libraries were prepared with the NEBNext Ultra DNA Library Prep Kit for Illumina (NEB) with a 500-bp insert size and sequencing was performed on a 2× 150bp Illumina NextSeq500 Medium at the VIB Nucleomics Core (Leuven, Belgium). Adapters and reads with a quality score below 20 were removed using BBduk2(*39*) with minlen=35 qtrim=rl trimq=20 hdist=1 tbo tpe and custom adapter reference file. Trimmed reads were aligned to *P. tricornutum* v2 assembly GCA_000150955.2 (*37*) (European Nucleotide Archive) using Burrows-Wheeler Alignment Tool (BWA) (*40*) algorithm BWA-MEM and processed and re-calibrated as described above. After the second round of recalibration, SNPs were called in three different ways:

*Germline SNP calling with GATK HaplotypeCaller a) joint genotyping with GATK 4.2.1 and b) for individual samples with GATK 3.7.0*: First, germline SNPs were called with HaplotypeCaller with either a) -ERC GVCF option and gVCF files were combined with CombineGVCFs and then jointly genotyped with GenotypeGVCFs b) or without the -ERC GVCF option and joint genotyping. SNPs were filtered with QD < 2.0, QUAL < 30.0, SOR > 3.0, FS > 60.2, MQ < 40.0, MQRankSum < -12.5, ReadPosRankSum < -8.0 and DP < 10 and indels with QD < 2.0, QUAL < 30.0, FS > 200.0, ReadPosRankSum < -20.0, DP < 10 using VariantFiltration and filtered SNPs and indels were removed with SelectVariants. These set of germline SNPs were used to build a Panel of Normals for following pairwise comparison of mother and daughter cell cultures using Mutect2 and for cross-verification of LOH regions identified in Mutect2.

*Pairwise comparison of mother and daughter cultures with GATK Mutect2:* All GATK algorithms were version 4.2.1 if not stated otherwise. First, SNPs were called on each sample with Mutect2 in tumor-only mode. Next, a vcf file for each sample was created by moving the sample-level AF allele-fraction annotation were moved into the INFO field for each sample using VariantsToTable. The Panel of Normals was prepared using the germline SNPs dataset for individual samples generated by HaplotypeCaller as a germline-resource by first calling the SNPs for each sample with Mutect2 in tumor-only mode, then merging all files with CombineVariants (GATK3.7.0) and finally creating the Panel of Normals with CreateSomaticPanelOfNormals. A Panel of Normals file for each of pairwise comparison was prepared by masking germline variants from the respective individual samples using germline SNPs called by HaplotypeCaller using CatVariants (GATK3.7.0) to merge SNPs from both samples and then masking them using SelectVariants with -XL option. Finally, LOH regions and *de novo* mutations were detected by Mutect2 SNP call with vcf file with sample-level AF allele-fraction in the INFO field used as --germline-resource, the respective masked Panel of Normals file used as –pon, --genotype-germline-sites true and either the mother culture used as tumor sample and daughter culture used as normal to detect LOH events in the daughter culture or vice-versa to detect *de novo* mutations in daughter culture (Tables S8 and Data S3). Called SNPs were filtered with FilterMutectCalls and filtered SNPs were removed with SelectVariants –excludeFiltered. As a control, the T1 with T2 time point of each mother culture were compared. A minimum of three consecutive SNPs missing in the daughter or T2 mother culture was considered as a LOH event and all LOH regions were reexamined in the datasets of germline SNPs obtained either by individual SNP calling or joint genotyping. The nature of the LOH was judged by comparison of coverage of the LOH region and its surrounding heterozygous borders and through Sanger resequencing of the LOH border (Table S8 and Data S3).

*CNV detection*: The copy-number variation was detected using GATK 4.1.7 CNV detection pipeline. First, intervals list with bin length set to 100 bp and -interval-merging-rule

OVERLAPPING_ONLY was prepared with PreprocessIntervals and collect raw counts were collected using CollectReadCounts and CNV panel of normals was generated by CreateReadCountPanelOfNormal with --minimum-interval-median-percentile 5.0 setting. Standardized copy ratios and denoised copy ratios were obtained using DenoiseReadCounts against panel of normals. Reference and alternative allele counts at common germline sites called on all samples by **HaplotypeCaller in GVCF** mode and jointly genotyped were obtained using **CollectAllelicCounts for each sample. Segments of contiguous copy ratios were acquired by ModelSegments in a paired analysis with --denoised-copy-ratios and --allelic-counts from the daughter culture and --normal-allelic-counts from the respective mother culture. Amplified, deleted and copy-neutral segments were called with CallCopyRatioSegments with default settings and plotted using** PlotModeledSegments. Detected CNV events were cross-verified in the datasets of germline SNPs obtained by SNP calling and joint genotyping and in estimated coverage counts per 10 bp obtained using bedtools 2.2.28 coverage. Further, identified LOH in tandem repeat on chromosome 5 in DC1.2 was verified by Sanger sequencing and duplication on chromosome 23 in DC1.3 was confirmed by qPCR quantification (see below).

***Re-sequencing of identified LOH events in mother versus daughter cell culture comparison and in CNV analysis:*** The nature of LOH events identified by pairwise comparison between mother and daughter cell culture was first judged by visual comparison of the sample coverage at the LOH region and the neighboring region. Next, the region was amplified by emulsion PCR as described in the Methods, PCR products were cloned into the pGEM-T vector and individual clones were sequenced. If the LOH region sequence was found together with both alleles of neighboring heterozygous SNP(s), the region was considered as copy-neutral LOH. If only a single allele was found and the coverage data corresponded, the region was considered as a deletion. Predicted heterozygous SNPs in regions between identified LOH events on chromosome 26 in DC2.1 culture were amplified by a standard PCR using Phusion High-Fidelity DNA Polymerases (Thermo Scientific) according to the manufacturers' instructions and sequenced by Sanger sequencing to verify their heterozygosity (Data S3). The same approach was used to verify the hetero/homozygosity of three SNPs on chromosome 27 that were predicted as LOH in culture DC2.2, but were called as homozygous by independent SNP calling also in the mother culture MC2. The LOH on chromosome 5 in DC1.2 was identified as CNV, but PCR amplification confirmed presence of two alleles with different length (differing in 1960 bp). The longer allele contained a tandemly duplicated region while in the short allele the duplication was missing. Subsequent Sanger sequencing of alleles showed that culture DC1.2 contains two short alleles with LOH of SNPs in the surrounding region (Fig. S6 and Data S3).

*qPCR quantification of duplication at chromosome 23 in DC1.4 culture:*

The relative copy number variation on chromosome 23 was examined in daughter cultures DC1.1, DC1.2 and DC1.3 in comparison with mother culture MC1 by quantitative real-time PCR (qPCR). DNA was extracted as described above and concentration was adjusted to 48 pg/μl for each sample. Two primer pairs were designed into the region containing putative duplication on chromosome 23 in DC1.3, two primer pairs were located on chromosome 23 outside the duplicated region and two primers pairs were targeting loci outside of chromosome 23, one on chromosome 6 and one on chromosome 22 (Fig. S6 and Data S3). qPCR was performed using the SYBR green kit (Roche) with 100 nM primers and 0.125 μl DNA in a total volume of 5 μl per reaction. qPCR amplification reactions were run and analyzed on the LightCycler 480 (Roche) with following cycling conditions: 10 min polymerase pre-incubation at 95°C and 45 cycles of amplification at 95°C for 10 s, 60°C for 15 s, and 72°C for 15 s. Melting curves were recorder after the last cycle by heating from 65 to 95°C. For each reaction, three technical repeats were performed. Data were analyzed using qbase+ (Biogazelle) (*58*) with copy number analysis option and with the mother culture MC1 as a reference sample and loci on distal arm of chromosome 23 (locus D) chromosome 6 (locus E) and chromosome 22 (locus F) as reference targets.

PtUMPS read-out system

*PtUMPS cultures*: Strains *ptumps-1bp* and *ptumps-1368bp* were generated based on a previously described protocol (*59*). Briefly, Cas9 ribonucleoprotein (RNP) complexes were assembled to target the *PtUMPS* locus, at either the gUMPS1 site or the gUMPS4 site (Data S3). An equimolar mixture of RNP gUMPS1 and RNP gUMPS4 (4 μg each) was bombarded into wild-type *P. tricornutum* Pt1 8.6 (CCMP2561) cells. Two rounds of selection were made on silicate-free F/2 medium (Sigma) plates supplemented with 50 μg/ml uracil (Sigma) and 300 μg/ml 5-fluoroorotic acid (5-FOA; ThermoFisher). Cell lysates were then prepared to serve as a template for genotyping. PCRs using the Q5 High Fidelity DNA polymerase (New England Biolabs) and primers UMPS_5UTR_F and UMPS_3UTR_R (Data S3) were performed to amplify the *PtUMPS* locus. The generated amplicons were subcloned employing the CloneJET PCR cloning kit (Thermo Scientific) and analyzed through Sanger sequencing. The *ptumps-320bp* strain was prepared and described previously under the name UA17 (*59*). *PtUMPS* mutant strains were maintained in conditions described above in 1xTMB medium supplemented with 50 μg/ml uracil and 100 μg/ml 5-FOA to prevent the restoration of wild-type allele.

*PtUMPS 14 day cultivation in non-selective conditions:* Cells densities of *ptumps-1bp, ptumps-320bp* and *ptumps-1368bp* were estimated using Bürker counting chamber and $20x10^6$ cells from were harvested by centrifugation at 1216 x g for 5 minutes. The cell pellet was washed four times with 50 ml of 1xTMB medium, then resuspended in 750 ml of 1x TMB medium supplemented with 50 μg/ml uracil and resulting cultures were cultivated in 12h/12h light/dark cycle. After 7 days, another 750 ml of fresh medium supplemented with uracil were added. After 14 days, cultures were harvested by centrifugation, cell density was estimated and $50x10^6$ cells from the culture were washed four times with TMB medium and plated on 1% agar ½ TMB medium 245 x 245 mm Nunc™ Square BioAssay plates (ThermoFisher) and incubated in 18h/6h light/dark cycle at 20°C for 6 weeks. Resulting colonies were manually counted (Table S11). Colonies selected for sequencing of *PtUMPS* locus were transferred to fresh 1xTMB medium and grown cell cultures were harvested as described above. The *PtUMPS* locus was amplified with primers PtUMPS-1 and PtUMPS-2 (Data S3) using two consecutive rounds of emulsion PCR. PCR products were cloned to pGEM-T vector and sequenced by Sanger sequencing.

<u>Estimation of interhomolog recombination frequency:</u>

~$50x10^6$ cells of three *ptumps-1bp* and five independent *ptumps-1368bp* cell subcultures started from a single cell were harvested by centrifugation at 1216 x g for 5 minutes. The cell pellet was washed four times with 50 ml of 1xTMB medium, then resuspended in 1xTMB medium. The cell density was determined and 2 to 6 replicas of either $25x10^6$ or $20x10^6$ cells were immediately plated on 1% agar ½ TMB medium and incubated as described above and incubated in 18h/6h light/dark cycle at 20°C for 6 weeks. Resulting colonies were manually counted (Table S12).

The frequency if interhomolog recombination was calculated based on the number of uracil prototrophic colonies after the immediate transfer of *ptumps-1368bp* strain from 5-FOA- and uracil-supplemented medium (only mutant cells survive) on plates without uracil (only cells that restored the wild-type allele survive). As the recombination that restored the wild-type *PtUMPS* allele had to occur within 1368-bp region, first the incidence of LOH events per 1000 bp was estimated for each replica by dividing the number of colonies by 1.38. The exact nuclear genome size of *P. tricornutum* was determined from genome fasta file using awk as 27,450,724 bp. Taking into account the genome length and number of plated cells, the recombination rate was recalculated per whole genome of 100 cells. As the copy-neutral LOH is detectable in half of the cases of mitotic interhomolog recombination due to random segregation of sister chromatids during mitotic metaphase, the number was multiplied by 2 to obtain the rate of mitotic recombination. Average value obtained from data for all replicas was $4.2 \times 10^{-2}$. The rate of reciprocal cross-overs on a 120-kb region in *S. cerevisiae* was estimated at $4 \times 10^{-5}$ per cell division and the rate of gene conversion events as $3.5 \times 10^{-6}$ per cell division (*60*) or $3.3 \times 10^{-3}$ per cell division for interstitial LOH, $1.4 \times 10^{-3}$ for terminal LOH genome-wide (*61*). The frequency of reciprocal cross-overs and gene conversion on a 120-kb region were combined and recalculated first per 1 kb and subsequently per 11.89 Mb *S. cerevisiae* genome. The resulting rate of interhomolog recombination in *S. cerevisiae* was calculated as $4.3 \times 10^{-3}$ in case of 120 kb region or $4.7 \times 10^{-3}$ in the case of the genome-wide studies.

Effect of treatment with cadmium, $H_2O_2$ and (E,E)-2,4-decadienal on interhomolog recombination at PtUMPS locus

The ranges of concentrations of chemicals used for treatments were first surveyed in literature, then selected concentrations were tested by treatment of Pt1 *P. tricornutum* strain for seven days. Afterwards, the cell survival of mock and treated cells was compared and the maximal dose that did not cause decrease in cell density was selected as the maximal dose in the respective experiment. *ptumps-1bp* and *ptumps-1368bp* cell cultures were harvested by centrifugation at 1216 x g  for 5 minutes and the cell pellet was washed four times with 50 ml of 1xTMB medium, then resuspended in 50 ml of 1xTMB medium. Cell density per ml was estimated and $25x10^6$ cells per replica were transferred to 200 ml of 1xTMB medium supplemented with 50 µg/ml uracil and respective treatment or mock treatment. For cadmium treatment, $CdCl_2$ (Sigma-Aldrich) was added to final $Cd^{2+}$ concentrations of 5 µg/L and 50 µg/L from 4000× stock solution. For $H_2O_2$ treatment, 30% solution of $H_2O_2$ (Merck) was added to a final concentration of 5 µM or 50 µM $H_2O_2$. For (E,E)-2,4-Decadienal treatment, 200 µL of DMSO was added to the mock treated cell cultures and (E,E)-2,4-Decadienal (Sigma-Aldrich) was added to 0.1 µM and 1 µM final concentration from 1000x and 100x concentrated stock solution respectively. Cultures were incubated for 24h in 12h/12h light/dark cycle at 20°C, under photosynthetic LED light with an intensity of 160 µmol photons $m^{-2}$ $s^{-1}$ and with 100 rpm shaking. Afterwards, all samples were harvested by centrifugation at 1216 x g  for 5 minutes  and cell pellets were washed four times with 50 ml of 1xTMB medium and plated on 1% agar ½ TMB medium as described above and incubated in 18h/6h light/dark cycle at 20°C for 6 weeks. Resulting colonies were manually counted (Tables S13-15).

### Supplementary Text

<u>Calculation of the error rate in self-corrected long-read sequencing data used for haplotype counting</u>

The long-read sequencing technology is known to have a higher error rate than the short-read sequencing technology (*62*). To improve the read quality, we generated circular consensus sequencing (CCS) reads for the PacBio datasets and used self-correction for both PacBio and MinION reads (detailed description is available in the Methods). The self-correction was preferred over correction by ILLUMINA reads, as the ILLUMINA reads were used for calling a SNP dataset and this could introduce bias in subsequent haplotype counting. The long-sequencing reads were further processed after alignment to the genome by removal of soft clipping, correction of CIGAR string and selection of uniquely mapped reads. The error rate in corrected and aligned PacBio and MinION files was estimated using Alfred (*63*) (Table S16).

The haplotype counting script processes only positions of reliable SNPs selected in ILLUMINA sequencing data and removes all reads containing insertions or deletions or other bases than the expected reference and alternative allele at the selected sites. Therefore, only the mismatch rate is relevant for the haplotype counting. Each position with a mismatch can result in one of the other three bases different from the reference. If the probability of mismatch to the reference is considered identical for each possible mismatched base, it is equal to one third. As only one of these bases is accepted by the haplotype counting script, the final probability of a mismatch at positions of reliable SNPs is equal to one third of the mismatch rate. Thus, probability of error at selected SNP sites used for haplotype determination ranged between

1.46% for *S. robusta* PacBio genome-wide sequencing and 0.05% for *P. tricornutum* T1 amplicon re-sequencing (Table S16).

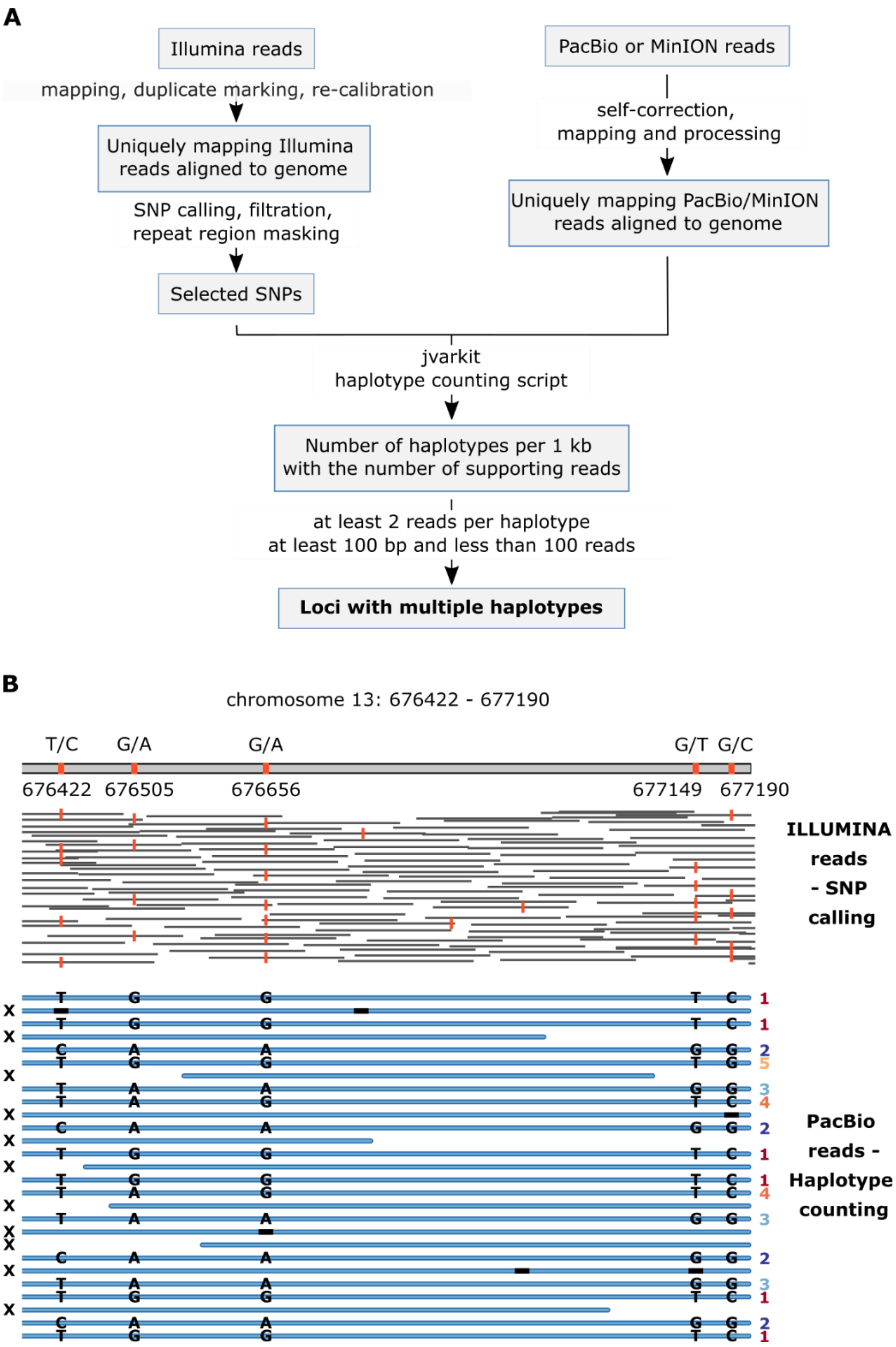


**Fig. S1. Schematic representation of genome-wide sequencing experiments and example.**

(**A**) Scheme of processing of publicly available genome-wide datasets for detection of loci with multiple haplotypes in *Seminavis robusta* and *Phaeodactylum tricornutum*. (**B**) Graphical scheme of one locus with multiple haplotypes detected in genome-wide haplotype counting in *P.*

*tricornutum*. X on the left denotes reads that were removed by the script, numbers on the right side denote the different haplotypes, black bar inside PacBio reads represents indels.

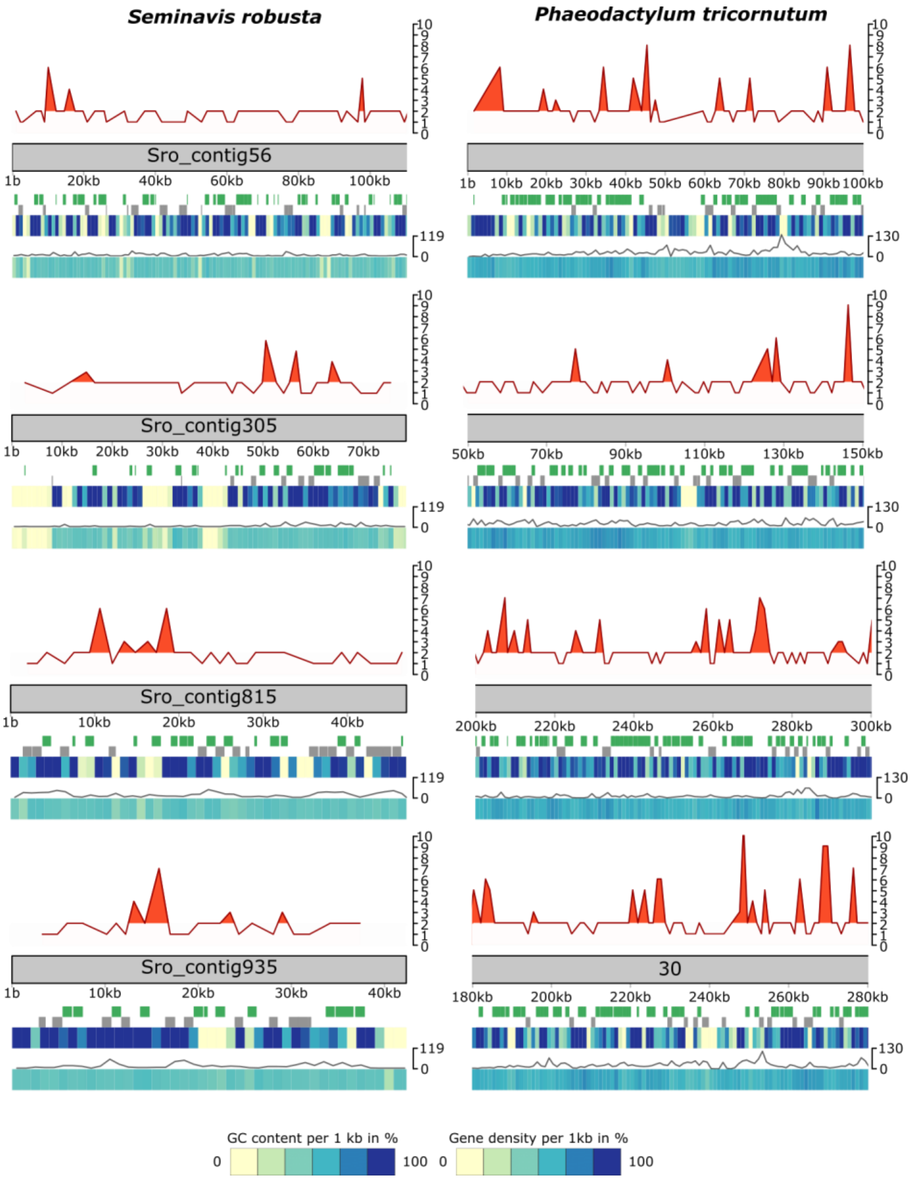


**Fig. S2. Randomly chosen examples of loci with multiple haplotypes.**

Above chromosome (in grey): number of found haplotypes (orange line), loci with more than two haplotypes (orange peaks). Below chromosome from top to bottom: loci with two haplotypes (green), loci with single haplotype (grey), gene density*, SNP density* (grey line), GC content*;

* per 1 kb.

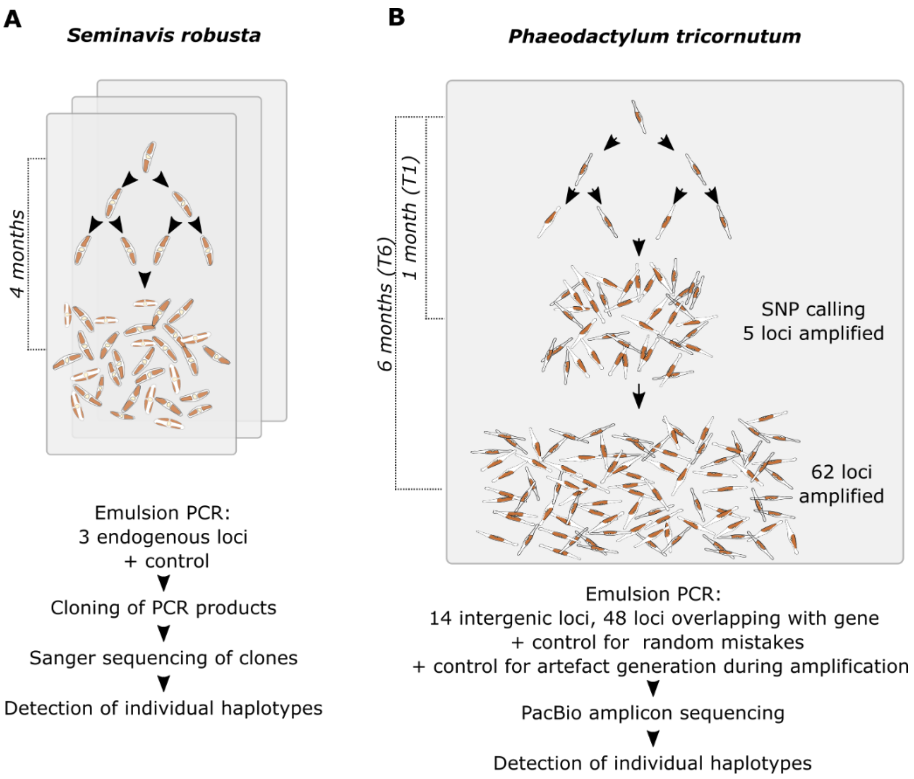


**Fig. S3. Graphical scheme of the mapping of haplotype accumulation in cultures freshly started from a single cell.**

Scheme of propagation of cultures from single cell in (A) *S. robusta* and (B) *P. tricornutum*.

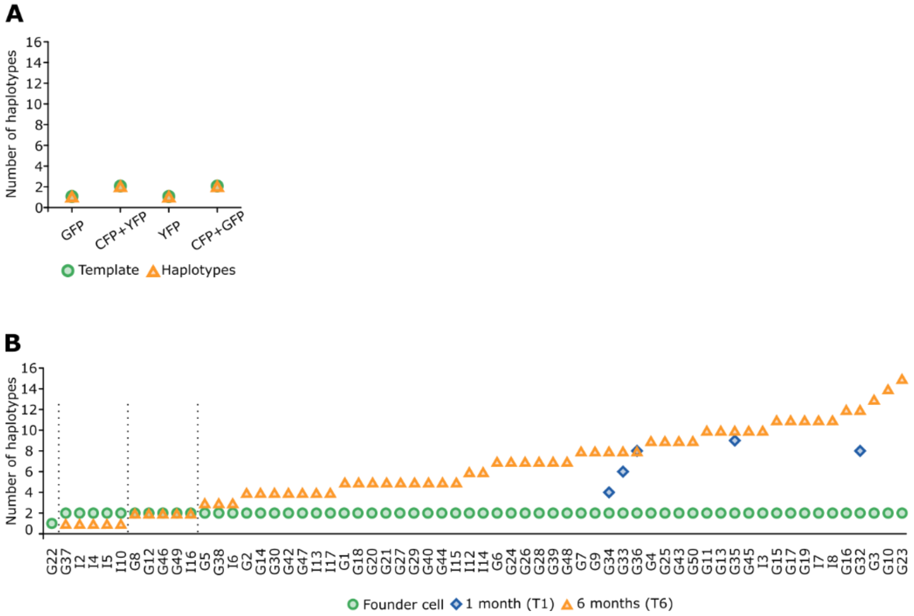


**Fig. S4. Detection of number of haplotypes in control reactions and at *P. tricornutum* endogenous loci at 1 and 6 months after cultivation from a single cell.**

(A) Control reactions (B) Haplotypes at all endogenous *P. tricornutum* loci detected 1 month (T1) and 6 months (T6) after single cell isolation.

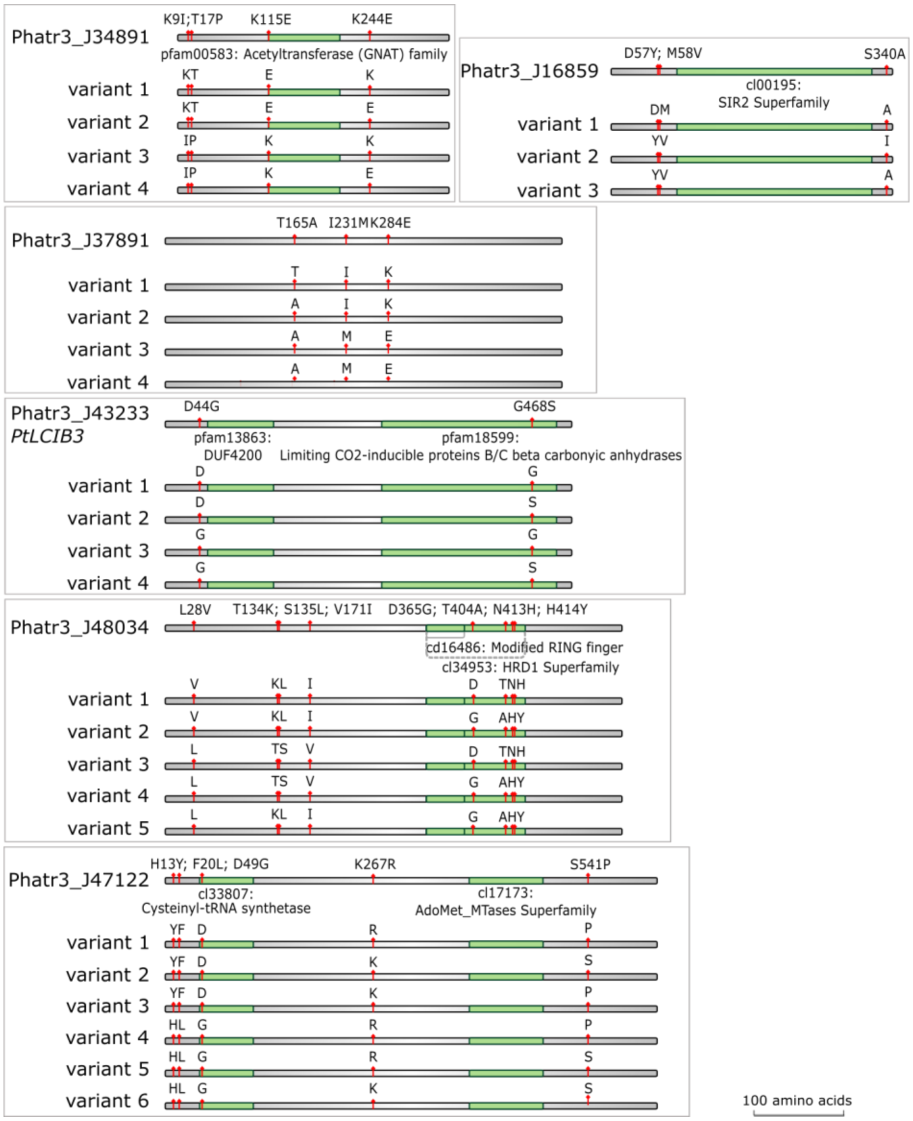


**Fig. S5. Schematic representation of predicted proteins variants resulting from haplotypes in genes fully covered by PacBio amplicon sequencing.**

Top line shows the position of amino acid variants on the protein indicated by red flags. Green regions depict conserved domains according to CDD/SPARKLE database (*64*). Lines below represent individual predicted variants.

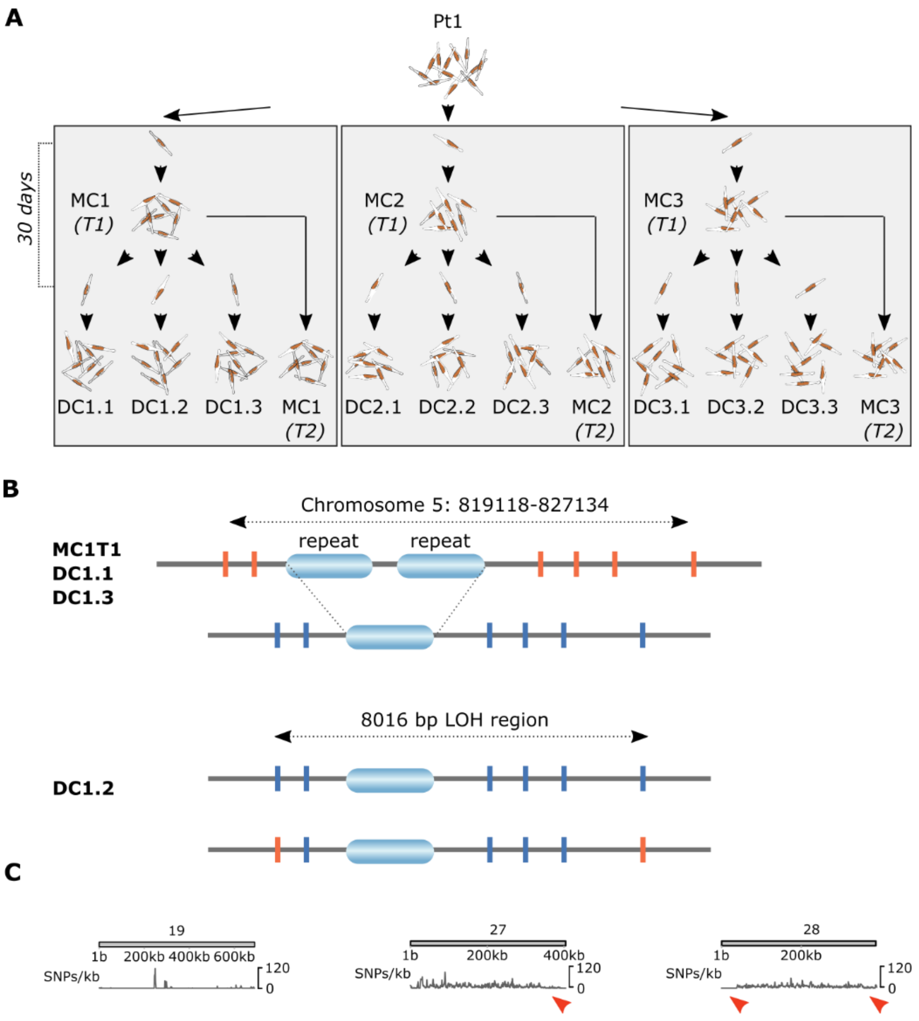


**Fig. S6. Scheme of the genome-wide genome rearrangements and LOH detection experiment and detected CNVs**

(**A**) Graphical scheme of the experiment. Three colonies from a single cell from the Pt1 culture were grown to obtain 3 mother cultures (MCx, T1 time point for deep sequencing). Thirty days after mother culture start, daughter cells DCx.1-DCx.3 were isolated from the respective mother cultures. (**B**) Scheme of copy-neutral LOH on chromosome 5 in DC1.2 identified in CNV analysis. Two alleles with different size were detected in the mother culture MC1 and sister cultures DC1.1

and DC1.3. Two alleles with identical size and SNPs were detected by Sanger sequencing in DC1.2. Horizontal grey bar - chromosome, vertical orange and blue bars - SNPs and blue ovals – region repeated in the longer allele. (**C**) Chromosomes with detected regions of low SNP density (pointed by red arrows on chromosome 27 and 28).

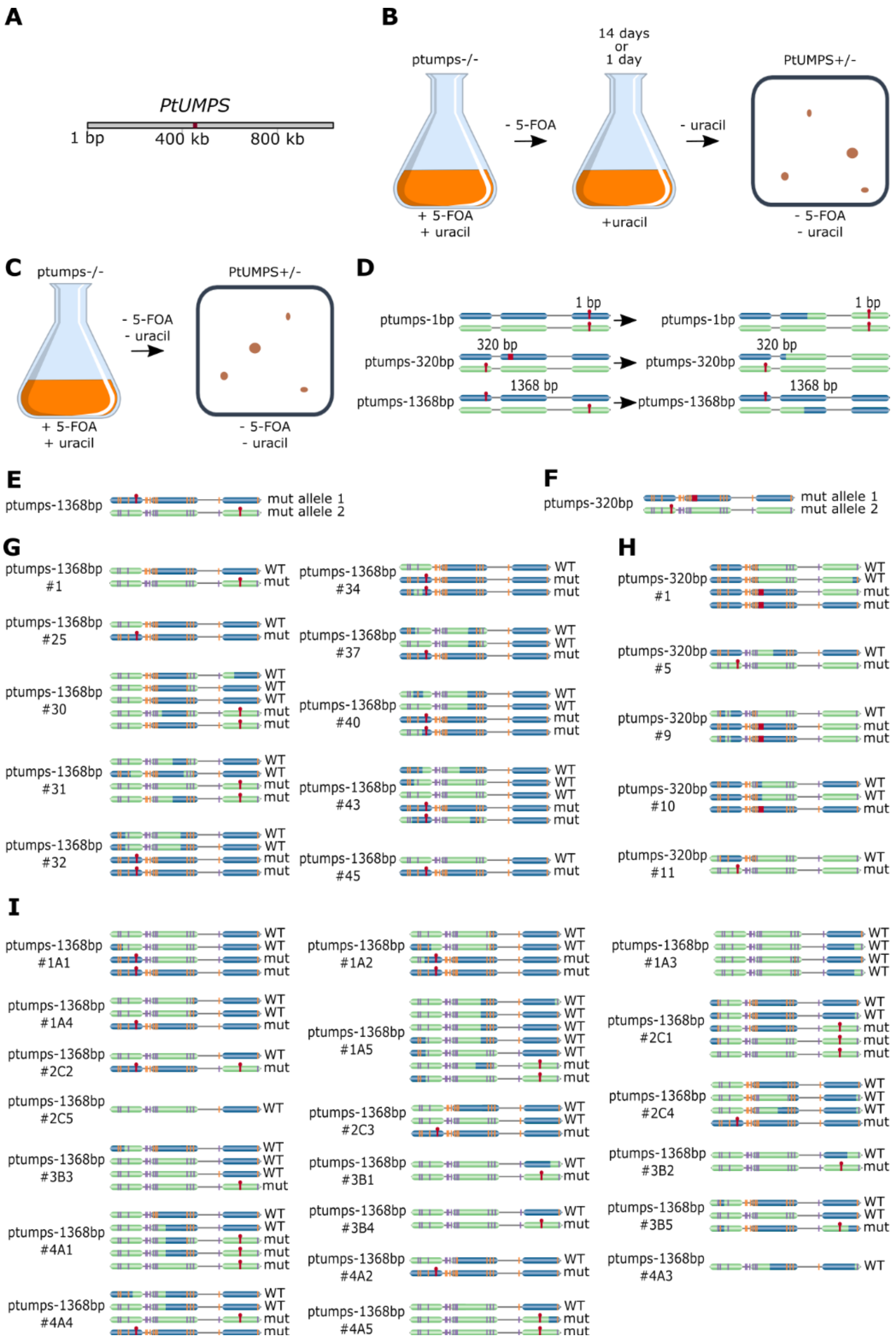


**Fig. S7. Graphical scheme and results of the *PtUMPS* system for detection of interhomolog recombination.**

(**A**) Position of the *PtUMPS* locus on chromosome 6. (**B**) Scheme of experiments allowing a period without selection pressure (+uracil) on *PtUMPS* locus. (**C**) Experimental design of interhomolog recombination frequency detection by immediate transfer from one selective condition (5-FOA and uracil; only *ptumps* mutants survive) to another (no uracil; only cells with WT *PtUMPS* allele survive). (**D**) Scheme of alleles of *PtUMPS* gene in strains used for detection of interhomolog recombination (left side) and hypothetical example of restoration of WT allele through interhomolog recombination in *ptumps-320bp* and *ptumps-1368bp*. Homologous chromosomes are depicted in blue and green, loss of function mutations are in red. (**E** to **F**) Position of silent SNPs (orange and mauve) and loss of function mutation (red) in (F) *ptumps-1368bp* and (E) *ptumps-320bp*. f, g, Alleles recovered in uracil prototrophic colonies of (**G**) *ptumps-1368bp*, (**H**) *ptumps-320bp* that were cultivated for 14 days in non-selective conditions and (**I**) after immediate transfer from 5-FOA and uracil containing medium to medium without uracil to select for cell undergoing interhomolog recombination within single round of cell division. WT – wild type; mut – mutant.

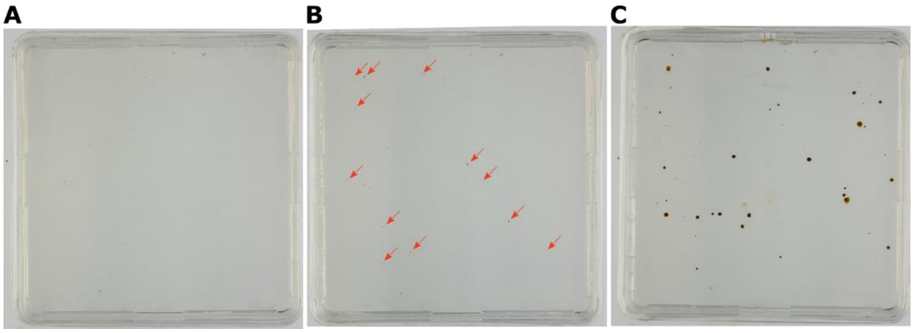


**Fig. S8. Uracil prototrophic colonies after cultivation of *PtUMPS* mutant strains in non-selective conditions for 14 days.**

(A) *ptumps-1bp*, (B) *ptumps-320bp* and (C) *ptumps-1368bp* strain

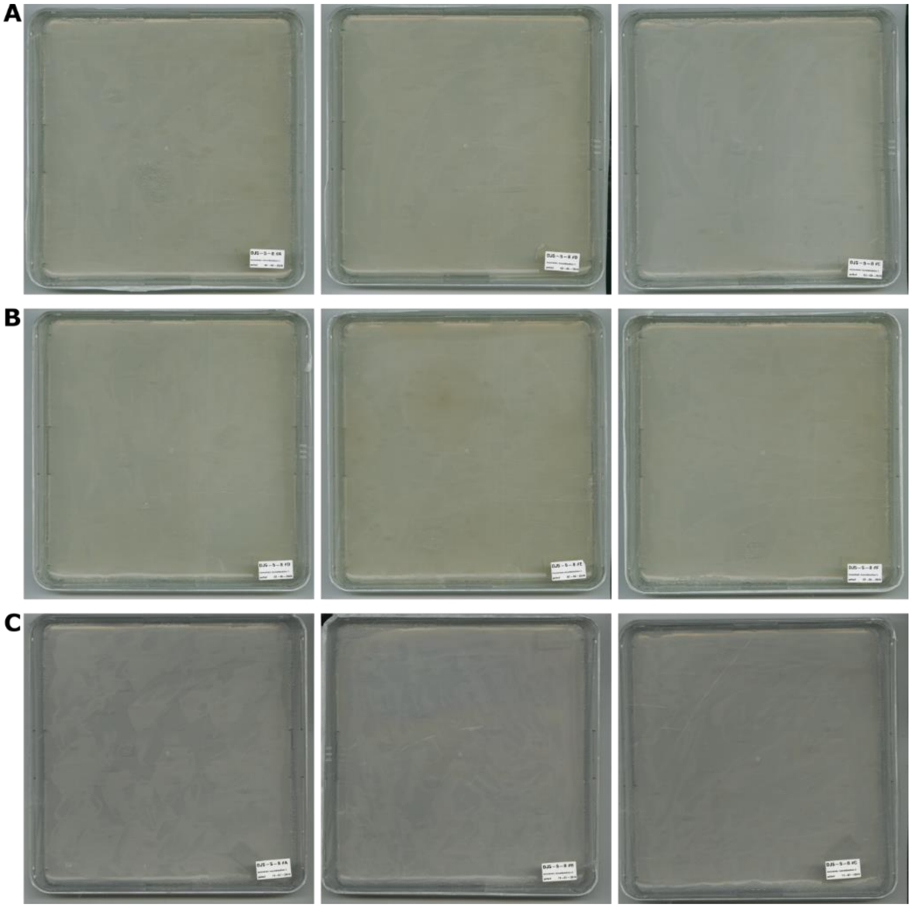

**Fig. S9. Estimation of interhomolog recombination rate in control *ptumps-1bp* strain**

No uracil prototrophic colonies were observed  after immediate transfer of *ptumps-1bp* PtUMPS mutant strains from 5-FOA and uracil supplemented medium to plates without uracil. (A) replica 1 (B) replica 2 (C) replica 3.

A

B

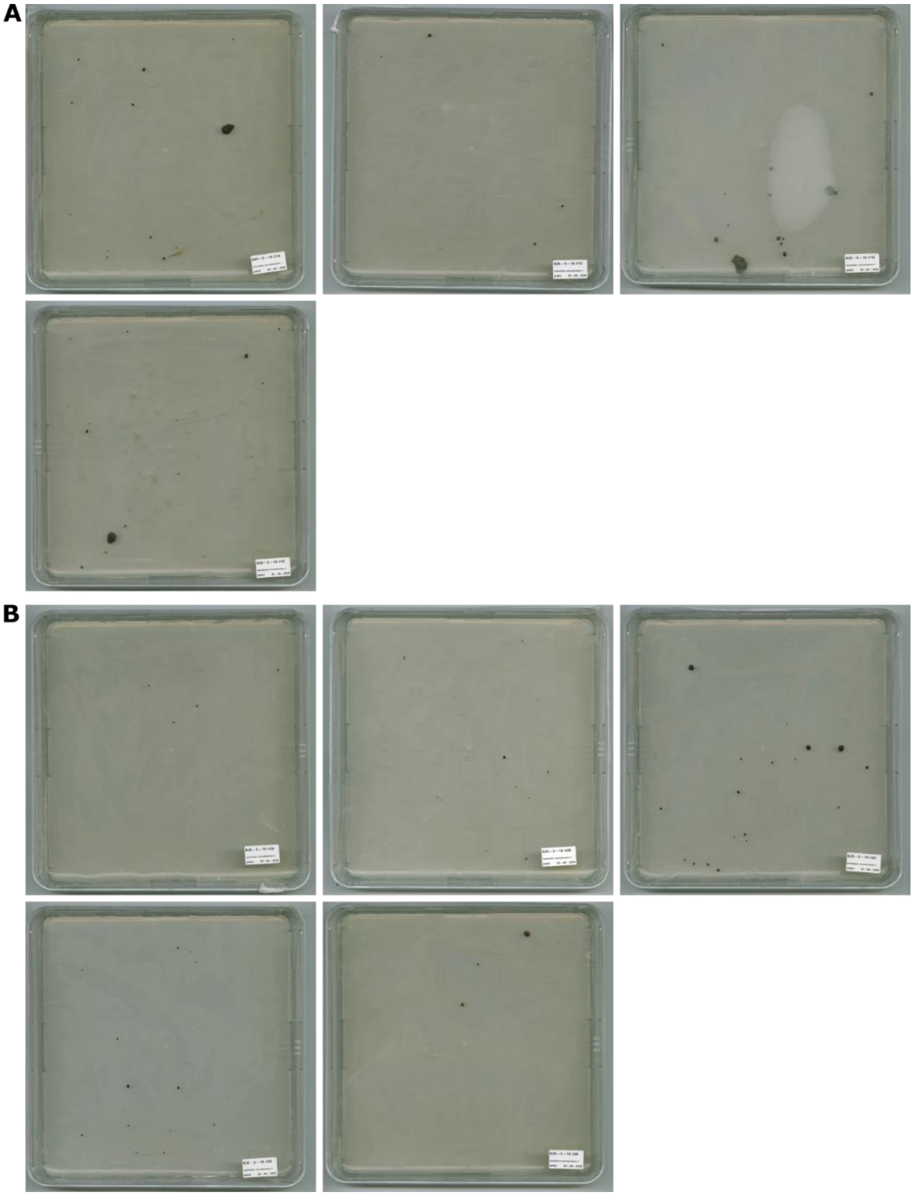

**Fig. S10. Estimation of interhomolog recombination rate in *ptumps-1368bp* strain, part1**

*ptumps-1368bp* uracil prototrophic colonies after immediate transfer of *PtUMPS* mutant strains from 5-FOA and uracil supplemented medium to plates without uracil to recover cells that were undergoing interhomolog recombination at *PtUMPS* locus and restored wild-type allele. (A) replica 1 (B) replica 2

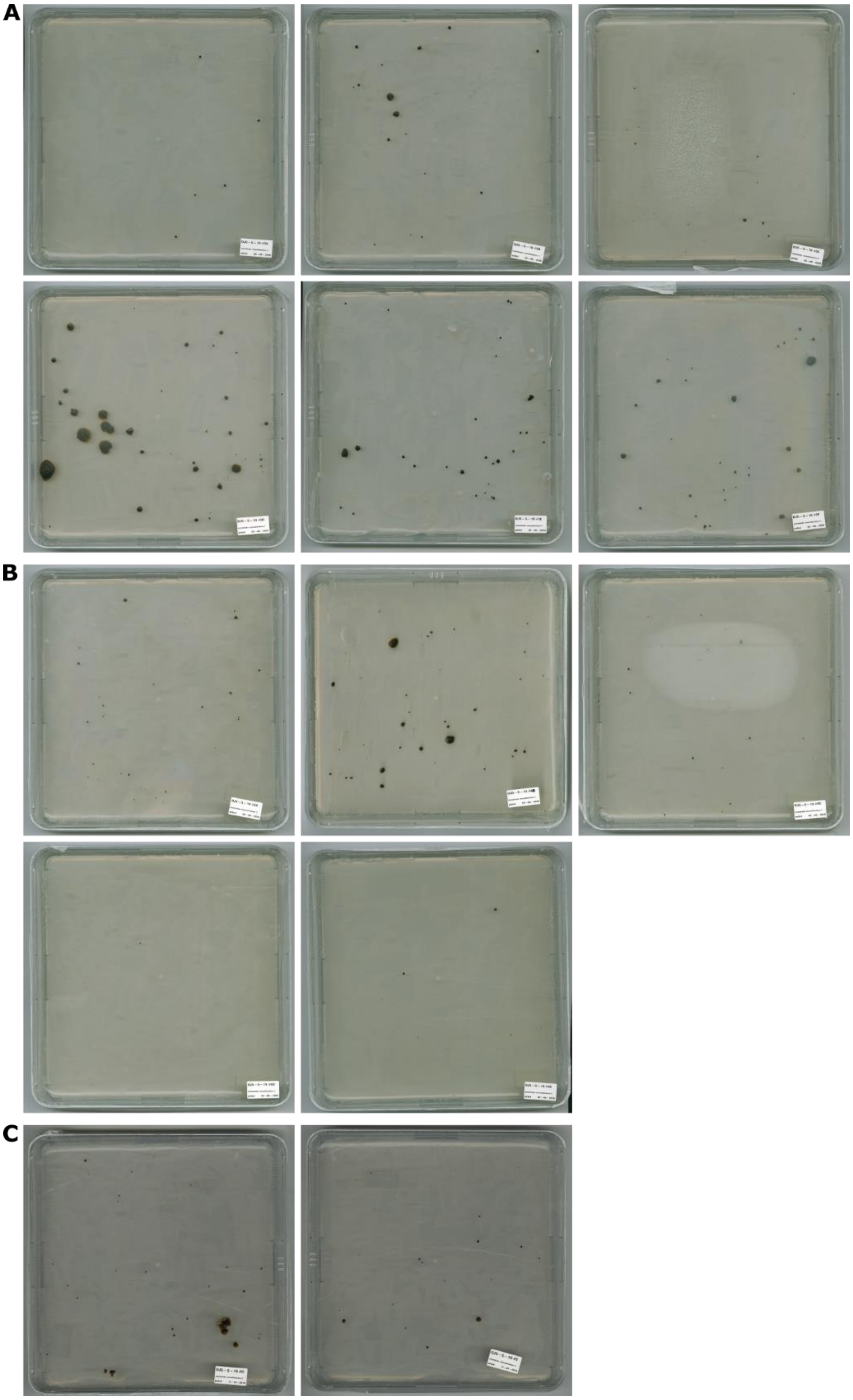

**Fig. S11. Estimation of interhomolog recombination rate in *ptumps-1368bp* strain, part 2**

*ptumps-1368bp* uracil prototrophic colonies after immediate transfer of *PtUMPS* mutant strains from 5-FOA and uracil supplemented medium to plates without uracil to recover cells that were undergoing interhomolog recombination at *PtUMPS* locus and restored wild-type allele. (A) replica 3 (B) replica 4, (C) replica 5

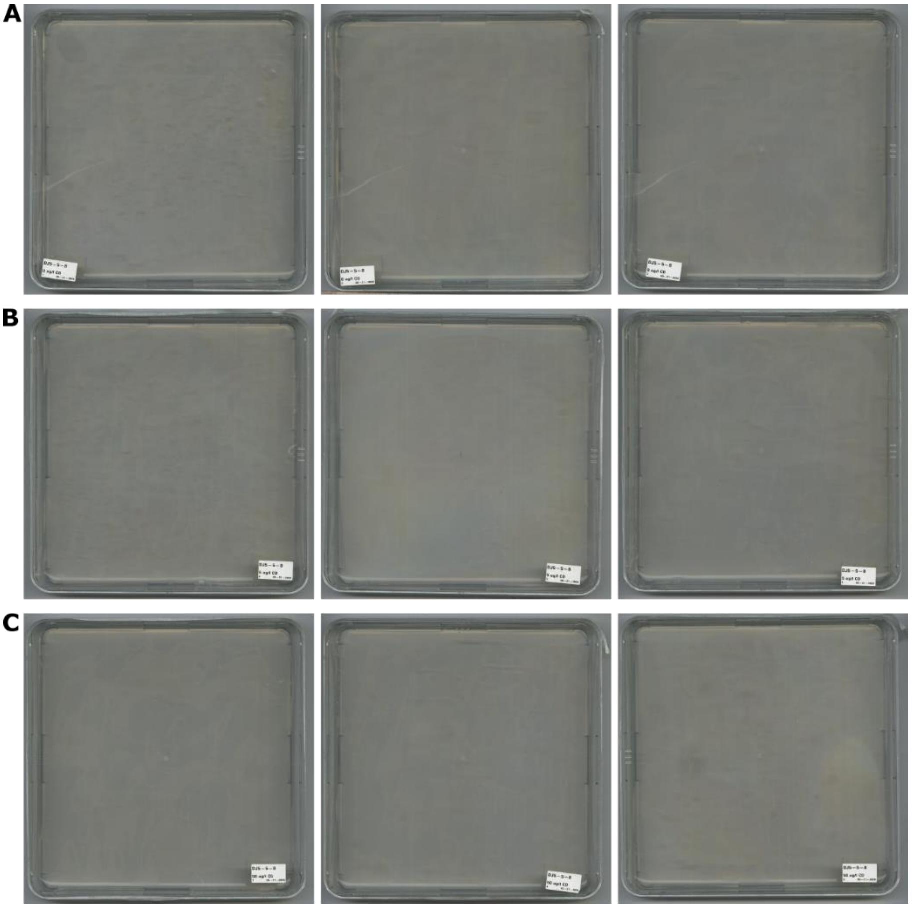


**Fig. S12. Uracil prototrophic colonies, corresponding to recombination events at *PtUMPS* locus, after exposure to stress induced by cadmium in control *ptumps-1bp* strain**

Plates for selection of uracil prototrophic colonies after cultivation of *ptumps-1bp* mutant strains in non-selective conditions for 1 day with or without cadmium treatment. (A) 0 μg/L cadmium (B) 5 μg/L cadmium and (C) 50 μg/L cadmium

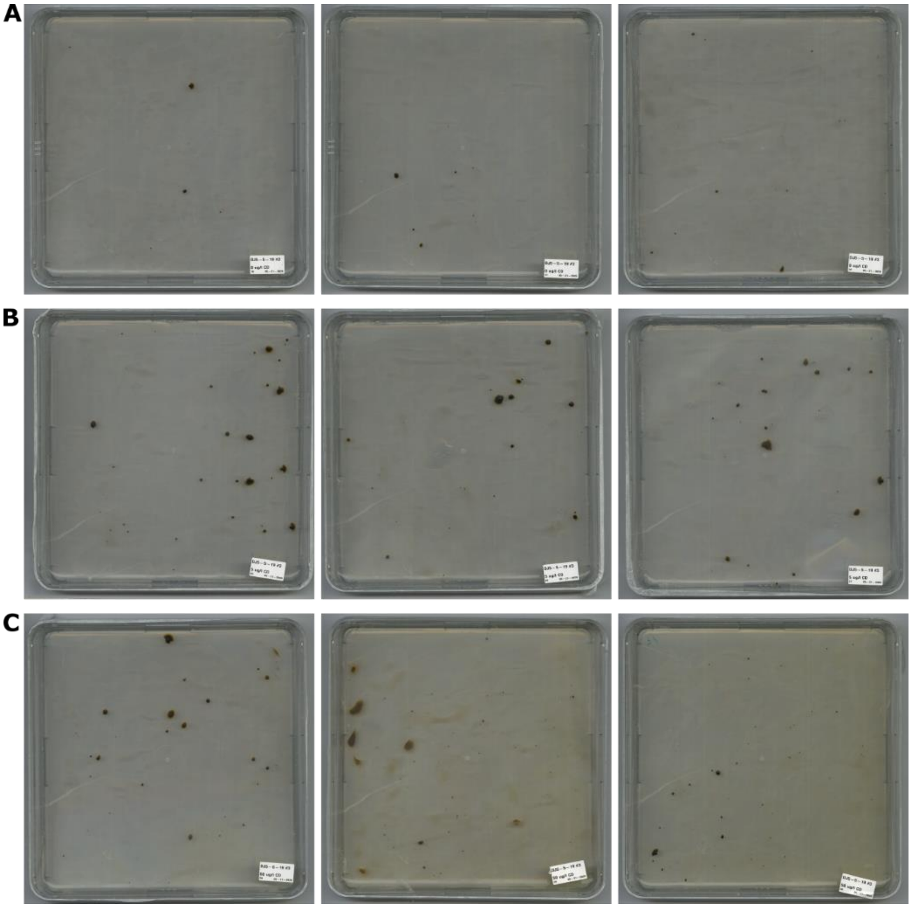

**Fig. S13. Uracil prototrophic colonies, corresponding to recombination events at *PtUMPS* locus, after exposure to stress induced by cadmium in *ptumps-1368bp* strain**

Plates for selection of uracil prototrophic colonies after cultivation of *ptumps-1368bp* mutant strains in non-selective conditions for 1 day with or without cadmium treatment. (A) 0 μg/L cadmium (B) 5 μg/L cadmium and (C) 50 μg/L cadmium

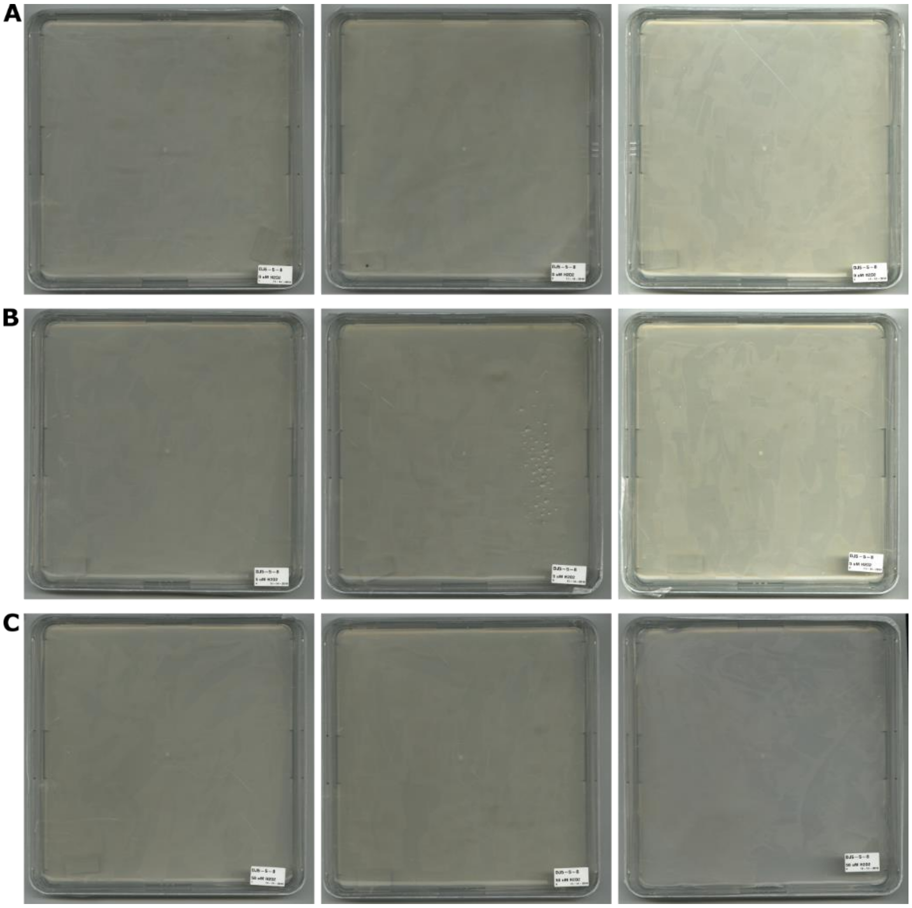

**Fig. S14. Uracil prototrophic colonies, corresponding to recombination events at *PtUMPS* locus, after exposure to stress induced by $H_2O_2$ in control *ptumps-1bp* strain**

Plates for selection of uracil prototrophic colonies after cultivation of *ptumps-1bp* mutant strains in non-selective conditions for 1 day with or without $H_2O_2$ treatment. (A) 0 µM $H_2O_2$ (B) 5 µM $H_2O_2$ and (C) 50 µM $H_2O_2$.

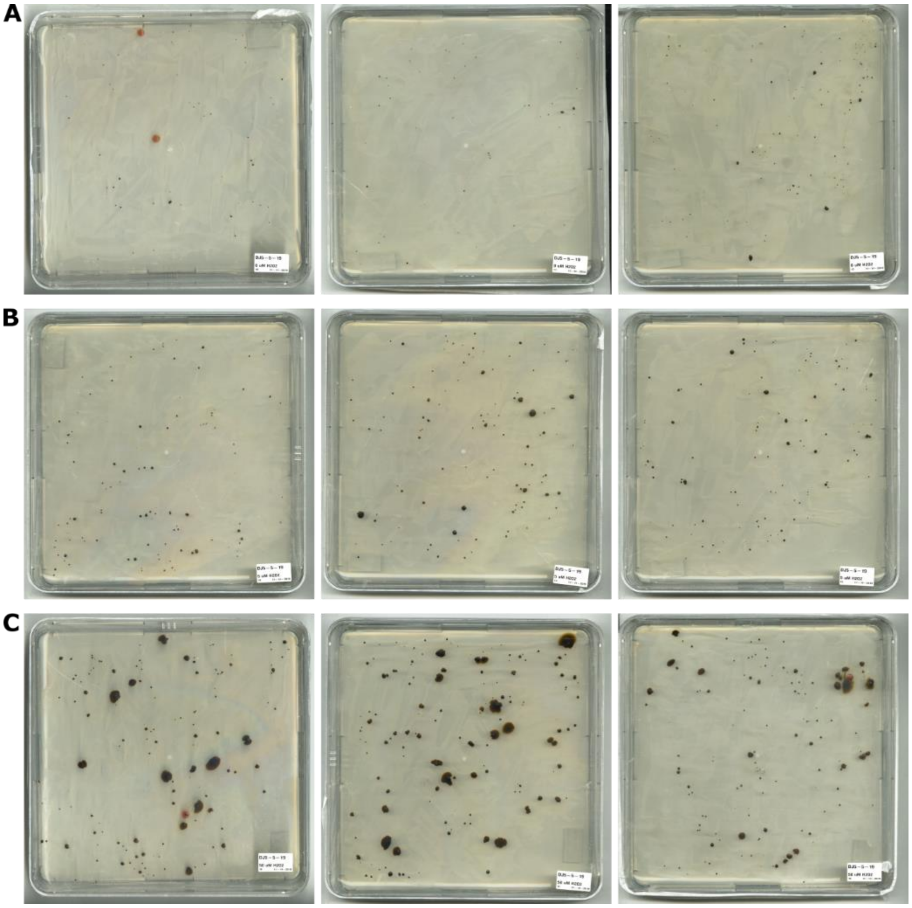

**Fig. S15. Uracil prototrophic colonies, corresponding to recombination events at *PtUMPS* locus, after exposure to stress induced by $H_2O_2$ in *ptumps-1368bp* strain**

Plates for selection of uracil prototrophic colonies after cultivation of *ptumps-1368bp* mutant strains in non-selective conditions for 1 day with or without $H_2O_2$ treatment. (A) 0 μM $H_2O_2$ (B) 5 μM $H_2O_2$ and (C) 50 μM $H_2O_2$.

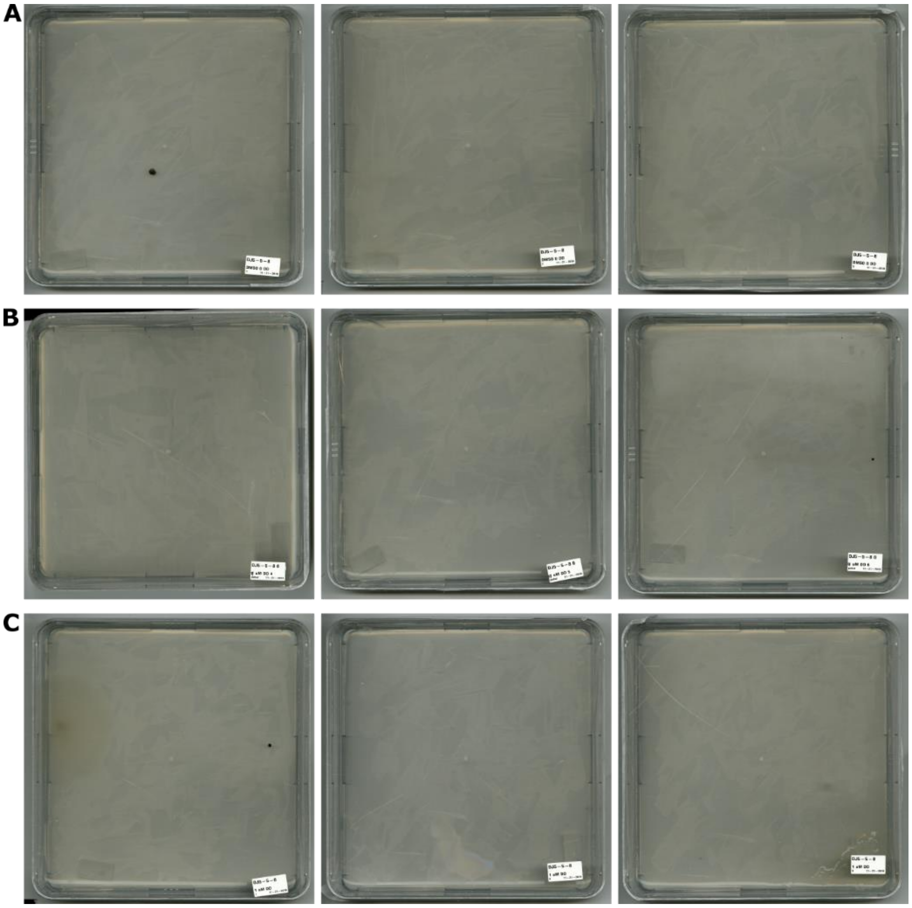

**Fig. S16. Uracil prototrophic colonies, corresponding to recombination events at *PtUMPS* locus, after exposure to stress induced by (E,E)-2,4-Decadienal in control *ptumps-1bp* strain**

Plates for selection of uracil prototrophic colonies after cultivation of *ptumps-1bp* mutant strains in non-selective conditions for 1 day with or without (E,E)-2,4-Decadienal treatment. (A) 0 μM (E,E)-2,4-Decadienal (B) 0.1 μM (E,E)-2,4-Decadienal and (C) 1 μM (E,E)-2,4-Decadienal

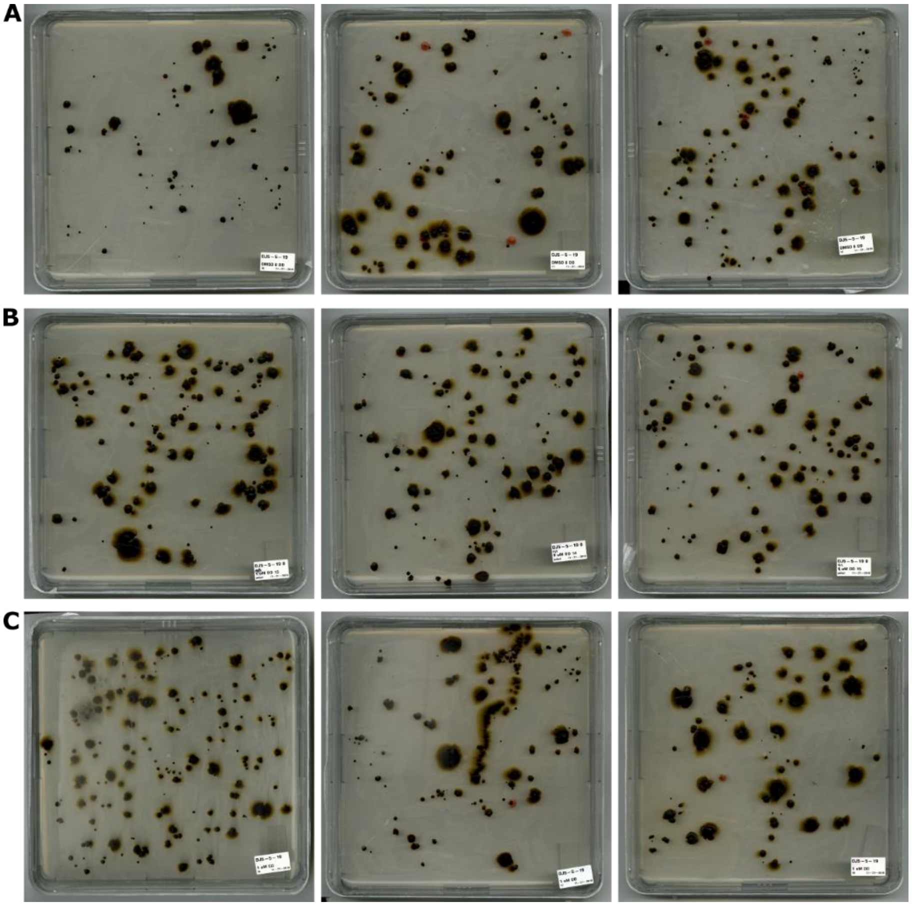


**Fig. S17. Uracil prototrophic colonies, corresponding to recombination events at *PtUMPS* locus, after exposure to stress induced by (E,E)-2,4-Decadienal in *ptumps-1368bp* strain**

Plates for selection of uracil prototrophic colonies after cultivation of *ptumps-1368bp* mutant strains in non-selective conditions for 1 day with or without (E,E)-2,4-Decadienal treatment. (A) 0 μM (E,E)-2,4-Decadienal (B) 0.1 μM (E,E)-2,4-Decadienal and (C) 1 μM (E,E)-2,4-Decadienal

**Table S1. Overview of genome-wide haplotype counting in diatoms *S. robusta* and *P. tricornutum*, yeast *Saccharomyces cerevisiae* and plant *Arabidopsis thaliana***

| **Species** | **Genome size** | **Coverage** | **Ploidy** | **Loci with multiple haplotypes** |
|---|---|---|---|---|
| *Seminavis robusta* | 125.5 Mb | ~14x | 2n | 1405 |
| *Phaeodactylum tricornutum* | 27.4 Mb | ~33x | 2n | 3380 |
| *Arabidopsis thaliana* Ler | 135 Mb | ~60.5 | 2n | 83 |
| *Saccharomyces cerevisiae* GLBRCY22-3 | 12 Mb | ~21.5 | 1n | 3 |

**Table S2. Characteristics of loci with multiple haplotypes found in *S. robusta* and *P. tricornutum***

| | Whole genome average | | Loci with multiple haplotypes | |
|---|---|---|---|---|
| | **Total number** | **Percent** | **Total number** | **Percent** |
| ***Seminavis robusta* (1405 loci)** | | | | |
| GC content | | 48.5 % | | 48.8 % |
| SNPs total | 489799 | 100 % | 7714 | 100% |
| SNPs in intergenic regions | 149782 | 30.58 % | 2662 | 34.50 % |
| SNPs in protein coding genes | 339890 | 69.39 % | 5050 | 65.47 % |
| Intron | 17300 | 3.53 % | 233 | 3.03 % |
| Exon | 322590 | 65.86 % | 4817 | 62.44 % |
| Functional RNAs | 127 | 0.03 % | 2 | 0.03 % |
| ***Phaeodactylum tricornutum* (3380 loci)** | | | | |
| GC content | | 48.77 % | | 49.04 % |
| SNPs total | 290164 | 100% | 22531 | 100% |
| SNPs in intergenic regions | 110727 | 38.16 % | 6767 | 30.03 % |
| SNPs in protein coding genes | 178906 | 61.65 % | 15735 | 69.83 % |
| Intron | 16271 | 5.61 % | 1242 | 5.51 % |
| Exon | 162635 | 56.04 % | 14493 | 64.32 % |
| Pseudogenes | 425 | 0.15 % | 27 | 0.12 % |
| Functional RNAs | 106 | 0.04 % | 2 | 0.01 % |

**Table S3. Verification of haplotype diversity in *S. robusta* at selected loci**

| Locus | Number of SNPs | Culture | Number of haplotypes at 4 months after single cell isolation | Number of supporting reads for each haplotype |
|---|---|---|---|---|
| Sro_contig211: 7509-8241 | | | | |
| | 11 | Sr1 | 5 | 26; 15; 1; 1; 1 |
| | 11 | Sr2 | 4 | 23; 18; 8; 1 |
| | 11 | Sr3 | 3 | 26; 25; 2 |
| Sro_contig2103: 8397-9162 | | | | |
| | 5 | Sr1 | 3 | 19; 12; 1 |
| | 5 | Sr2 | 3 | 8; 6; 1 |
| | 5 | Sr3 | 2 | 24; 13 |
| Sro_contig872: 16034-16975 | | | | |
| | 4 | Sr1 | 3 | 24; 16; 1 |
| | 4 | Sr2 | 5 | 19; 14; 1; 1; 1 |
| | 4 | Sr3 | 6 | 29; 19; 3; 2; 1; 1 |
| Sro_contig556:54453-55487 - control mix of plasmids containing two alleles of the locus | | | | |
| | 3 | - | 2 | 44; 19 |

**Table S4. The number of haplotypes at control amplicons obtained by PacBio amplicon sequencing. Only haplotypes supported by at least 1% of reads from the total read count or at least 2 reads were added to the haplotype count.**

| **Locus name** | **Barcode** | **Number of haplotypes** | **Coverage** | **List of reads supporting individual haplotypes** |
|---|---|---:|---|---|
| GFP | 1 | 1 | 26771 | 26756 |
| YFP | 2 | 1 | 8853 | 8848 |
| CFP+YFP | 1 | 2 | 7131 | 4361, 2724 |
| CFP+GFP | 2 | 2 | 148 | 114, 34 |

**Table S5. The number of haplotypes of *P. tricornutum* loci used for the verification of haplotype diversity by PacBio amplicon sequencing, six months after single-cell isolation. Only haplotypes supported by at least 1% of reads from the total read count or at least 2 reads were added to the haplotype count.**

| **Locus name** | **Chrom** | **Start** | **End** | **Number of haplotypes** | **Coverage** | **List of reads supporting individual haplotypes** |
|---|---|---|---|---|---|---|
| G1 | 1 | 1003970 | 1005992 | 5 | 767 | 387, 126, 104, 9, 9 |
| G2 | 1 | 1002273 | 1004299 | 4 | 855 | 343, 287, 19, 13 |
| G3 | 20 | 379968 | 382045 | 13 | 3861 | 1228, 1135, 94, 92, 87, 79, 70, 70, 70, 58, 50, 42, 42 |
| G4 | 21 | 456389 | 458435 | 9 | 2754 | 919, 604, 79, 73, 57, 46, 44, 41, 38 |
| G5 | 23 | 16577 | 18629 | 3 | 3528 | 2990, 88, 66 |
| G6 | 24 | 409922 | 411940 | 7 | 2934 | 770, 764, 92, 40, 40, 35, 33 |
| G7 | 25 | 457868 | 459883 | 8 | 2659 | 1170, 166, 52, 39, 39, 38, 34, 30 |
| G8 | 26 | 45670 | 47665 | 2 | 803 | 758, 17 |
| G9 | 27 | 254779 | 256783 | 8 | 1399 | 831, 26, 25, 24, 19, 18, 14, 14 |

| | | | | | | |
|---|---|---|---|---|---|---|
| G10 | 28 | 312712 | 314788 | 14 | 1419 | 470, 452, 52, 51, 36, 36, 29, 23, 18, 17, 16, 15, 15, 15 |
| G11 | 2 | 365525 | 367572 | 10 | 1341 | 354, 353, 30, 28, 23, 23, 16, 16, 15, 14 |
| G12 | 30 | 122641 | 124669 | 2 | 2704 | 2140, 240 |
| G13 | 33 | 8238 | 10281 | 10 | 4128 | 1138, 1120, 89, 80, 60, 56, 51, 51, 50, 48 |
| G14 | 4 | 1263120 | 1265150 | 4 | 3344 | 1097, 537, 500, 48 |
| G15 | 10 | 282249 | 284312 | 11 | 3493 | 1180, 1144, 153, 142, 142, 127, 92, 85, 39, 37, 36 |
| G16 | 11 | 401452 | 403473 | 12 | 1999 | 583, 560, 31, 29, 28, 27, 25, 23, 22, 21, 20,20 |
| G17 | 12 | 643173 | 645247 | 11 | 366 | 156, 12, 9, 8, 5, 5, 4, 4, 4, 4, 4 |
| G18 | 13 | 676389 | 678462 | 5 | 3393 | 1162, 938, 65, 53, 38 |
| G19 | 14 | 334406 | 336466 | 11 | 3151 | 1136, 1092, 96, 67, 67, 49, 49, 36, 36, 35, 33 |
| G20 | 15 | 485547 | 487569 | 5 | 605 | 269, 221, 10, 8, 8 |
| G21 | 16 | 563372 | 565452 | 5 | 5667 | 1736, 1687, 112, 64, 61 |
| G22 | 19 | 110953 | 112967 | 1 | 6349 | 6345 |

| | | | | | | |
|---|---|---|---|---|---|---|
| G23 | 17 | 573662 | 575737 | 15 | 1210 | 384, 328, 44, 34, 29, 26, 25, 23, 21, 19, 19, 19, 18, 15, 13 |
| G24 | 17 | 127422 | 129499 | 7 | 2038 | 684, 518, 59, 32, 29, 26, 26 |
| G25 | 18 | 43706 | 45792 | 9 | 1276 | 486, 352, 39, 38, 27, 18, 17, 14, 14 |
| G26 | 18 | 418603 | 420622 | 7 | 526 | 344, 18, 11, 9, 9, 7, 7 |
| G27 | 18 | 421123 | 423150 | 5 | 1110 | 450, 289, 28, 25, 13 |
| G28 | 4 | 987941 | 990015 | 7 | 417 | 160, 155, 12, 7, 6, 5, 5 |
| G29 | 6 | 601781 | 603775 | 5 | 401 | 147, 145, 8, 7, 5 |
| G30 | 6 | 603744 | 605766 | 4 | 568 | 230, 223, 12, 11 |
| G32 | 13 | 103145 | 105183 | 12 | 878 | 310, 264, 28, 19, 19, 16, 16, 13, 10, 10, 10, 9 |
| G33 | 27 | 205731 | 207770 | 8 | 988 | 340, 299, 26, 21, 18, 14, 11, 11 |
| G34 | 20 | 101923 | 103985 | 8 | 654 | 316, 116, 14, 12, 9, 8, 7, 7 |
| G35 | 12 | 519921 | 521994 | 10 | 706 | 268, 225, 19, 17, 17, 13, 12, 11, 9, 8 |
| G36 | 2 | 961633 | 963692 | 8 | 688 | 246, 208, 34, 17, 13, 12, 12, 11 |
| G37 | 11 | 393793 | 395845 | 1 | 727 | 692 |
| G38 | 4 | 1021280 | 1023325 | 3 | 841 | 303, 255, 10 |

| | | | | | | |
|---|---|---|---|---|---|---|
| G39 | 18 | 541891 | 543937 | 7 | 833 | 339, 328, 61, 41, 23, 20, 9 |
| G40 | 15 | 770178 | 772260 | 5 | 772 | 269, 239, 11, 9, 9 |
| G42 | 13 | 150502 | 152542 | 4 | 687 | 342, 217, 11, 10 |
| G43 | 8 | 525255 | 527273 | 9 | 913 | 401, 319, 25, 16, 16, 14, 12, 12, 10 |
| G44 | 6 | 18873 | 20900 | 5 | 1377 | 571, 435, 41, 15, 14 |
| G45 | 1 | 904526 | 906542 | 10 | 1473 | 650, 539, 47, 44, 36, 28, 22, 19, 18, 18 |
| G46 | 3 | 730840 | 732851 | 2 | 994 | 923, 11 |
| G47 | 29 | 264288 | 266345 | 4 | 1332 | 579, 505, 23, 16 |
| G48 | 1 | 2400393 | 2402421 | 7 | 1860 | 858, 634, 31, 27, 25, 24, 22 |
| G49 | 1 | 1198020 | 1200061 | 2 | 1282 | 494, 494 |
| G50 | 1 | 388317 | 390345 | 9 | 1476 | 615, 298, 263, 35, 30, 29, 21, 19, 17 |
| I2 | 1 | 716566 | 718619 | 1 | 2283 | 2279 |
| I3 | 9 | 932002 | 934038 | 10 | 1109 | 359, 278, 36, 32, 31, 19, 18, 14, 14, 12 |
| I4 | 5 | 27292 | 29302 | 1 | 1125 | 1122 |
| I5 | 8 | 350011 | 352014 | 1 | 1801 | 1787 |
| I6 | 4 | 195053 | 197089 | 3 | 1613 | 655, 536, 19 |
| I7 | 9 | 937756 | 939779 | 11 | 486 | 152, 145, 21, 14, 11, 11, 9, 8, 6, 5, 5 |

| | | | | | | |
|---|---|---|---|---|---|---|
| I8 | 10 | 919965 | 921984 | 11 | 395 | 251, 11, 10, 10, 8, 7, 4, 4, 4, 4, 4 |
| I10 | 12 | 191252 | 193312 | 1 | 538 | 538 |
| I12 | 2 | 1412376 | 1414425 | 6 | 1126 | 523, 326, 28, 23, 16, 14 |
| I13 | 2 | 1086792 | 1088839 | 4 | 1426 | 681, 624, 56, 46 |
| I14 | 2 | 1098502 | 1100525 | 6 | 979 | 440, 350, 25, 19, 18, 14 |
| I15 | 30 | 104972 | 107013 | 5 | 965 | 509, 256, 12, 10, 10 |
| I16 | 33 | 44228 | 46283 | 2 | 690 | 576, 11 |
| I17 | 6 | 57110 | 59115 | 4 | 1718 | 679, 613, 22, 19 |

**Table S6. Change in number of recovered haplotypes over time in *P. tricornutum***

| Locus name | Coordinates | Number of SNPs | Samples harvested at 1 month after single cell isolation | | Samples harvested at 6 month after single cell isolation | |
|---|---|---|---|---|---|---|
| | | | Number of haplotypes | Coverage | Number of haplotypes | Coverage |
| G32 | 13:103145-105183 | 10 | 8 | 3903 | 12 | 878 |
| G33 | 27:205731-207770 | 18 | 6 | 5712 | 8 | 988 |
| G34 | 20:101923-103985 | 28 | 4 | 3140 | 8 | 654 |
| G35 | 12:519921-521994 | 12 | 9 | 2992 | 10 | 706 |
| G36 | 2:961633-963692 | 12 | 8 | 5225 | 8 | 688 |

**Table S7. Genes fully covered in PacBio amplicon sequencing**

| Protein ID | Locus name | Locus coordinates | Number of haplotypes at T6 | Number of predicted protein isoforms |
|---|---|---|---|---|
| Phatr3_J48737 | G22 | 19: 110953-112967 | 1 | 1 |
| Phatr3_J46763 | G37 | 11: 393793-395845 | 1 | 1 |
| Phatr3_J47158 | G17 | 12: 643173-645247 | 11 | 1 |
| Phatr3_J44026 | G46 | 3: 730840- 732851 | 2 | 1 |
| Phatr3_J40830 | G8 | 26: 45670-47665 | 2 | 2 |
| Phatr3_EG01422 | G44 | 6: 18873-20900 | 5 | 2 |
| Phatr3_J9020 | G49 | 1: 1198020-1200061 | 2 | 2 |
| Phatr3_J3062 | G50 | 1: 388317-390345 | 9 | 2 |
| Phatr3_J8717 | G45 | 1:904526-906542 | 10 | 2 |
| Phatr3_J42757 | G1 | 1: 1003970-1005992 | 5 | 2 |
| Phatr3_J38628 | G21 | 16 563372-565452 | 5 | 2 |
| Phatr3_J38754 | G24 | 17:127422-129499 | 7 | 2 |
| Phatr3_J38755 | G24 | 17:127422-129499 | 7 | 2 |
| Phatr3_J38927 | G23 | 17: 573662-575737 | 15 | 2 |
| Phatr3_J16859 | G47 | 29:264288-266345 | 4 | 3 |
| Phatr3_J34891 | G29 | 6:601781-603775 | 5 | 4 |
| Phatr3_J43233 | G48 | 1:2400393-2402421 | 7 | 4 |
| Phatr3_J37891 | G19 | 14:334406-336466 | 11 | 4 |
| Phatr3_J48034 | G40 | 15:770178-772260 | 5 | 5 |
| Phatr3_J47122 | G35 | 12:519921-521994 | 10 | 6 |

**Table S8. Regions identified by pairwise comparison of mother and daughter cultures in the genome-wide LOH and CNV analysis**

| Sample | Region and number of SNPs in pairwise comparison of mother and daughter cells | Confirmation by Sanger sequencing or qPCR | Region and covered SNPs in GATK joint genotyping |
|---|---|---|---|
| **DC1.2** | 5:819118-827134, detected as CNV | Copy-neutral LOH in a region involving a tandem duplication (Sanger sequencing) | 5:819118-827134, 73 SNPs |
| **DC1.3** | 2:4818-5114, 3 SNPs | Copy-neutral LOH (Sanger sequencing) | 2:4818-5114 3 SNPs |
| **DC1.3** | 17:16150-17794, 8 SNPs | Copy-neutral LOH (Sanger sequencing) | 17:15681-17978 19 SNPs |
| **DC1.3** | 23:7901-39400, detected as CNV | Duplication (qPCR) | 23:7901-39400, 474 SNPs |
| **DC1.3** | 24:7360-7974, 3 SNPs | Copy-neutral LOH (Sanger sequencing) | 24:7360-7809 3 SNPs |
| **DC2.1** | 26:19535-176510, 9 SNPs | Confirmed as single deletion (Sanger sequencing) | 26: 19535 – 177391, SNPs: 2018 |
| **DC2.1** | 26:27495-69347, 58 SNPs | | |
| **DC2.1** | 26:74508-75499, 3 SNPs | | |
| **DC2.1** | 26:109659-133578, 43 SNPs | | |
| **DC2.1** | 26:139651-158982, 45 SNPs | | |

| | | | |
|---|---|---|---|
| **DC2.1** | 26:162893-176510, 16 SNPs | | |
| **DC2.2** | 27:373856-387563, 3 SNPs | Not confirmed | Called as homozygous in all MC2 and DC2 samples |
| **DC3.1** | 23:8373-39308, 57 SNPs | Deletion (Sanger sequencing) | 23:8070-38628 475 SNPs |

**Table S9. Gene covered by duplication on chromosome 23 in DC1.3**

| **Locus** | **Gene Identifier** | **Protein ID** | **Gene name (if available)** |
|---|---|---|---|
| 23 : 6743-8328 | ptri27310 | Phatr3_J49547 | |
| 23 : 8871-10156 | ptri27330 | Phatr3_J40282 | |
| 23 : 10378-12190 | ptri27350 | Phatr3_J30446 | |
| 23 : 13563-15056 | ptri27390 | Phatr3_J55067 | |
| 23 : 15368-17272 | ptri27410 | Phatr3_J49550 | |
| 23 : 17726-19318 | ptri27430 | Phatr3_J49551 | |
| 23 : 19840-20493 | ptri27450 | Phatr3_J16155 | |
| 23 : 21334-22059 | ptri27470 | Phatr3_J49552 | |
| 23 : 22329-24011 | ptri27490 | Phatr3_J49553 | |
| 23 : 24669-25199 | ptri27510 | Phatr3_J23324 | |
| 23 : 25949-29740 | ptri27530 | Phatr3_J49555 | |
| 23 : 30937-32655 | ptri27550 | Phatr3_J49556 | |
| 23 : 32879-34165 | ptri27570 | Phatr3_J49557 | Pt_HSF4.a |
| 23 : 35105-37350 | ptri27590 | Phatr3_EG01932 | Pt_HSF4.6b |

**Table S10. SNPs with high effect on protein function fixed by LOH and deletions**

| Culture | SNP position | Alleles | Fixed allele | Affected protein | Effect on protein |
|---|---|---|---|---|---|
| **DC3.1** | **23: 30947** | **G/A** | **A** | **Phatr3_J49556** | **W4X; fixation of premature stop codon** |
| DC3.1 | 23: 33129 | T/ TCAAAG | T | Phatr3_J49557 | Frameshift compensating mutations |
| DC3.1 | 23: 33130 | T/ TGTCTATCTTCATC | T | | |
| DC2.1 | 26: 28021 | TG/T | TG | Phatr3_EG01904 | Frameshift compensating mutations |
| DC2.1 | 26 :28023 | CGTCGT/T | CGTCGT | | |
| DC2.1 | 26: 32779 | A/ATTTC | ATTTC | Phatr3_J3656 | Frameshift compensating mutations |
| DC2.1 | 26: 32780 | AGGAG/A | A | | |
| DC2.1 | 26: 46285 | G/A | G | Phatr3_J40830 | W7X; fixation of functional allele |
| **DC2.1** | **26: 46961** | **A/T** | **T** | **Phatr3_J40830** | **K183X; fixation of premature stop codon** |

| | | | | | |
|---|---|---|---|---|---|
| DC2.1 | 26: 47348 | T/G | G | Phatr3_J40830 | X312E; removes stop codon, 3a.a. longer protein |
| DC2.1 | 26: 48144 | C/CT | C | Phatr3_J40831 | C393V,induces frameshift; fixation of functional allele |
| DC2.1<br>DC2.1 | 26: 56724<br>26: 56727 | ATC/A<br>A/ATG | A<br>ATG | Phatr3_J49989 | Frameshift compensating mutations |
| DC2.1 | 26: 140915 | T/A | T | Phatr3_J16517 | X360Y-protein 15a.a longer in alternative allele; fixation of original allele |
| DC2.1<br>DC2.1 | 26: 154678<br>26: 154681 | TGG/T<br>A/AAC | TGG<br>A | Phatr3_EG00171 | Frameshift; fixation of functional allele |
| DC2.1 | 26: 163666 | TC/T | TC | Phatr3_J40874 | Frameshift; fixation of functional allele |
| **DC2.1** | **26: 163668** | **C/CG** | **CG** | **Phatr3_J40874** | **Frameshift; fixation of premature stop codon, 70 a.a. shorter protein** |

**Table S11. Numbers of *ptumps-1bp, ptumps-320bp* and *ptumps-1368bp* uracil prototrophic colonies after a 14-day cultivation in non-selective conditions**

| **Strain** | **Number of plated cells** | **Number of uracil prototrophic colonies** |
|---|---|---|
| *ptumps-1bp* | $50 \times 10^6$ | 0 |
| *ptumps-320bp* | $50 \times 10^6$ | 12 |
| *ptumps-1368bp* | $50 \times 10^6$ | 83 |

**Table S12. Numbers of *ptumps-1bp* and *ptumps-1368bp* uracil prototrophic colonies after immediate transfer from 5-FOA- and uracil-supplemented medium to medium without uracil**

| **Strain** | **Subculture started from single cell** | **Replica** | **Number of plated cells** | **Number of uracil prototrophic colonies** |
|---|---|---|---|---|
| *ptumps-1bp* | #1 | 1 | $20 \times 10^6$ | 0 |
| *ptumps-1bp* | #1 | 2 | $20 \times 10^6$ | 0 |
| *ptumps-1bp* | #1 | 3 | $20 \times 10^6$ | 0 |
| *ptumps-1bp* | #2 | 1 | $20 \times 10^6$ | 0 |
| *ptumps-1bp* | #2 | 2 | $20 \times 10^6$ | 0 |
| *ptumps-1bp* | #2 | 3 | $20 \times 10^6$ | 0 |
| *ptumps-1bp* | #3 | 1 | $25 \times 10^6$ | 0 |
| *ptumps-1bp* | #3 | 2 | $25 \times 10^6$ | 0 |
| *ptumps-1bp* | #3 | 3 | $25 \times 10^6$ | 0 |
| *ptumps-1368bp* | #1 | 1 | $20 \times 10^6$ | 12 |
| *ptumps-1368bp* | #1 | 2 | $20 \times 10^6$ | 10 |
| *ptumps-1368bp* | #1 | 3 | $20 \times 10^6$ | 18 |
| *ptumps-1368bp* | #1 | 4 | $20 \times 10^6$ | 22 |
| *ptumps-1368bp* | #2 | 1 | $20 \times 10^6$ | 5 |
| *ptumps-1368bp* | #2 | 2 | $20 \times 10^6$ | 12 |
| *ptumps-1368bp* | #2 | 3 | $20 \times 10^6$ | 21 |

| | | | | |
|---|---|---|---|---|
| *ptumps-1368bp* | #2 | 4 | $20 \times 10^6$ | 16 |
| *ptumps-1368bp* | #2 | 5 | $20 \times 10^6$ | 13 |
| *ptumps-1368bp* | #3 | 1 | $20 \times 10^6$ | 6 |
| *ptumps-1368bp* | #3 | 2 | $20 \times 10^6$ | 17 |
| *ptumps-1368bp* | #3 | 3 | $20 \times 10^6$ | 27 |
| *ptumps-1368bp* | #3 | 4 | $20 \times 10^6$ | 48 |
| *ptumps-1368bp* | #3 | 5 | $20 \times 10^6$ | 29 |
| *ptumps-1368bp* | #3 | 6 | $20 \times 10^6$ | 32 |
| *ptumps-1368bp* | #4 | 1 | $20 \times 10^6$ | 31 |
| *ptumps-1368bp* | #4 | 2 | $20 \times 10^6$ | 51 |
| *ptumps-1368bp* | #4 | 3 | $20 \times 10^6$ | 19 |
| *ptumps-1368bp* | #4 | 4 | $20 \times 10^6$ | 6 |
| *ptumps-1368bp* | #4 | 5 | $20 \times 10^6$ | 7 |
| *ptumps-1368bp* | #5 | 1 | $25 \times 10^6$ | 33 |
| *ptumps-1368bp* | #5 | 2 | $25 \times 10^6$ | 21 |

**Table S13. Numbers of *ptumps-1bp* and *ptumps-1368bp* uracil prototrophic colonies after 24-h cadmium treatment in non-selective conditions**

| **Strain** | **Number of cells before plating** | **Replica** | **Treatment** | **Number of uracil prototrophic colonies** |
|---|---|---|---|---|
| *ptumps-1bp* | $25 \times 10^6$ | 1 | 0 µg/L | 0 |
| *ptumps-1bp* | $25 \times 10^6$ | 2 | 0 µg/L | 0 |
| *ptumps-1bp* | $25 \times 10^6$ | 3 | 0 µg/L | 0 |
| *ptumps-1bp* | $25 \times 10^6$ | 1 | 5 µg/L | 0 |
| *ptumps-1bp* | $25 \times 10^6$ | 2 | 5 µg/L | 0 |
| *ptumps-1bp* | $25 \times 10^6$ | 3 | 5 µg/L | 0 |
| *ptumps-1bp* | $25 \times 10^6$ | 1 | 50 µg/L | 0 |
| *ptumps-1bp* | $25 \times 10^6$ | 2 | 50 µg/L | 0 |
| *ptumps-1bp* | $25 \times 10^6$ | 3 | 50 µg/L | 0 |
| *ptumps-1368bp* | $25 \times 10^6$ | 1 | 0 µg/L | 12 |
| *ptumps-1368bp* | $25 \times 10^6$ | 2 | 0 µg/L | 13 |
| *ptumps-1368bp* | $25 \times 10^6$ | 3 | 0 µg/L | 19 |
| *ptumps-1368bp* | $25 \times 10^6$ | 1 | 5 µg/L | 39 |
| *ptumps-1368bp* | $25 \times 10^6$ | 2 | 5 µg/L | 34 |
| *ptumps-1368bp* | $25 \times 10^6$ | 3 | 5 µg/L | 40 |
| *ptumps-1368bp* | $25 \times 10^6$ | 1 | 50 µg/L | 41 |
| *ptumps-1368bp* | $25 \times 10^6$ | 2 | 50 µg/L | 92 |
| *ptumps-1368bp* | $25 \times 10^6$ | 3 | 50 µg/L | 54 |

**Table S14. Numbers of *ptumps-1bp* and *ptumps-1368bp* uracil prototrophic colonies after 24-h $H_2O_2$ treatment in non-selective conditions**

| **Strain** | **Number of cells before plating** | **Replica** | **Concentration of H2O2** | **Number of uracil prototrophic colonies** |
|---|---|---|---|---|
| *ptumps-1bp* | 25 x $10^6$ | 1 | **0 µM** | **1** |
| *ptumps-1bp* | 25 x $10^6$ | 2 | **0 µM** | **1** |
| *ptumps-1bp* | 25 x $10^6$ | 3 | **0 µM** | **0** |
| *ptumps-1bp* | 25 x $10^6$ | 1 | **5 µM** | **0** |
| *ptumps-1bp* | 25 x $10^6$ | 2 | **5 µM** | **0** |
| *ptumps-1bp* | 25 x $10^6$ | 3 | **5 µM** | **0** |
| *ptumps-1bp* | 25 x $10^6$ | 1 | **50 µM** | **0** |
| *ptumps-1bp* | 25 x $10^6$ | 2 | **50 µM** | **0** |
| *ptumps-1bp* | 25 x $10^6$ | 3 | **50 µM** | **0** |
| *ptumps-1368bp* | 25 x $10^6$ | 1 | **0 µM** | **54** |
| *ptumps-1368bp* | 25 x $10^6$ | 2 | **0 µM** | **52** |
| *ptumps-1368bp* | 25 x $10^6$ | 3 | **0 µM** | **93** |
| *ptumps-1368bp* | 25 x $10^6$ | 1 | **5 µM** | **117** |
| *ptumps-1368bp* | 25 x $10^6$ | 2 | **5 µM** | **114** |
| *ptumps-1368bp* | 25 x $10^6$ | 3 | **5 µM** | **123** |
| *ptumps-1368bp* | 25 x $10^6$ | 1 | **50 µM** | **143** |
| *ptumps-1368bp* | 25 x $10^6$ | 2 | **50 µM** | **149** |
| *ptumps-1368bp* | 25 x $10^6$ | 3 | **50 µM** | **141** |

**Table S15. Numbers of *ptumps-1bp* and *ptumps-1368bp* uracil prototrophic colonies after 24-h (E,E)-2,4-Decadienal treatment in non-selective conditions**

| Strain | Number of cells before plating | Replica | Concentration of (E,E)-2,4-Decadienal | Number of uracil prototrophic colonies |
|---|---|---|---|---|
| *ptumps-1bp* | $25 \times 10^6$ | 1 | 0 μM | **1** |
| *ptumps-1bp* | $25 \times 10^6$ | 2 | 0 μM | **0** |
| *ptumps-1bp* | $25 \times 10^6$ | 3 | 0 μM | **0** |
| *ptumps-1bp* | $25 \times 10^6$ | 1 | 0.1 μM | **0** |
| *ptumps-1bp* | $25 \times 10^6$ | 2 | 0.1 μM | **0** |
| *ptumps-1bp* | $25 \times 10^6$ | 3 | 0.1 μM | **1** |
| *ptumps-1bp* | $25 \times 10^6$ | 1 | 1 μM | **1** |
| *ptumps-1bp* | $25 \times 10^6$ | 2 | 1 μM | **0** |
| *ptumps-1bp* | $25 \times 10^6$ | 3 | 1 μM | **0** |
| *ptumps-1368bp* | $25 \times 10^6$ | 1 | 0 μM | **72** |
| *ptumps-1368bp* | $25 \times 10^6$ | 2 | 0 μM | **87** |
| *ptumps-1368bp* | $25 \times 10^6$ | 3 | 0 μM | **110** |
| *ptumps-1368bp* | $25 \times 10^6$ | 1 | 0.1 μM | **122** |
| *ptumps-1368bp* | $25 \times 10^6$ | 2 | 0.1 μM | **110** |
| *ptumps-1368bp* | $25 \times 10^6$ | 3 | 0.1 μM | **117** |
| *ptumps-1368bp* | $25 \times 10^6$ | 1 | 1 μM | **159** |
| *ptumps-1368bp* | $25 \times 10^6$ | 2 | 1 μM | **128** |
| *ptumps-1368bp* | $25 \times 10^6$ | 3 | 1 μM | **76** |

**Table S16. Error rate in processed long-read sequencing data used for haplotype counting estimated by Alfred.**

| | Error rate | Deletion rate | Insertion rate | Mismatch rate | Final probability of error at selected sites |
|---|---|---|---|---|---|
| *S. robusta* genome-wide PacBio sequencing | | | | | |
| | 5.23% | 0.54% | 0.27% | 4.42% | **1.46%** |
| *P. tricornutum* genome-wide MinION sequencing | | | | | |
| | 4.92% | 2.13% | 0.99% | 1.79% | **0.59%** |
| *P. tricornutum* PacBio amplicon re-sequencing T1 | | | | | |
| | 0.80% | 0.08% | 0.56% | 0.16% | **0.05%** |
| *P. tricornutum* PacBio amplicon re-sequencing T6 | | | | | |
| | 0.94% | 0.09% | 0.53% | 0,32% | **0.11%** |

**Data S1. (separate file) Details of the SNV positions and densities in the core genes.**

This table lists the (A) SNV positions in the exons present in the core genes, shown for the surface metagenome (10% departure from consensus, at least 20X coverage in the samples). (B) SNV positions in the exons present in the core genes, shown for the DCM metagenome (10% departure from consensus, at least 20X coverage in the samples). (C) SNV densities (%) of the exons present in the core genes. (D) SNV densities (%) of the core genes.

**Data S2. (separate file) List of loci with more than two haplotypes detected genome-wide in in profiled organisms**

This table lists the position, number of SNPs, number of total aligned reads, number of reads validated for haplotype counting, number of haplotypes and number of supporting reads for each of the loci with more than two expected haplotypes uncovered genome-wide haplotype analysis in *S. robusta*, *P. tricornutum*, *A. thaliana* and *S. cerevisiae*.

**Data S3. (separate file) List of material used throughout the study**

This file comprises information on diatom datasets and strains, list of loci and list of positions of SNPs present in control amplicons in re-sequencing of haplotype diversity in *S. robusta* and *P. tricornutum,* list of oligonucleotides used throughout the study and list of position of de novo SNPs found in comparison of mother and daughter cell cultures.

**References**


31. T. Mock *et al.*, Evolutionary genomics of the cold-adapted diatom Fragilariopsis cylindrus. *Nature* **541**, 536-540 (2017).
32. B. Langmead, S. L. Salzberg, Fast gapped-read alignment with Bowtie 2. *Nat Methods* **9**, 357-359 (2012).
33. H. Li *et al.*, The Sequence Alignment/Map format and SAMtools. *Bioinformatics* **25**, 2078-2079 (2009).
34. A. M. Eren *et al.*, Anvi'o: an advanced analysis and visualization platformfor 'omics data. *Peerj* **3**, (2015).
35. T. O. Delmont *et al.*, Single-amino acid variants reveal evolutionary processes that shape the biogeography of a global SAR11 subclade. *Elife* **8**, (2019).
36. C. M. Osuna-Cruz *et al.*, The Seminavis robusta genome provides insights into the evolutionary adaptations of benthic diatoms. *Nat Commun* **11**, 3320 (2020).
37. C. Bowler *et al.*, The Phaeodactylum genome reveals the evolutionary history of diatom genomes. *Nature* **456**, 239-244 (2008).
38. G. A. Van der Auwera *et al.*, From FastQ data to high confidence variant calls: the Genome Analysis Toolkit best practices pipeline. *Curr Protoc Bioinformatics* **43**, 11 10 11-11 10 33 (2013).
39. B. Bushnell, BBMap. *sourceforge.net/projects/bbmap/*.
40. H. Li, R. Durbin, Fast and accurate short read alignment with Burrows-Wheeler transform. *Bioinformatics* **25**, 1754-1760 (2009).
41. B. Institute, Picard Tools. *Broad Institute, GitHub repository*, (Accessed: 2019, version 2.6.0).
42. R. Poplin *et al.*, Scaling accurate genetic variant discovery to tens of thousands of samples. 201178 (2018).
43. A. Smit, Hubley, R., RepeatModeler Open-1.0. <*http://www.repeatmasker.org*>. (2008-2015).
44. A. Smit, Hubley, R & Green, P, RepeatMasker Open-4.0. <*http://www.repeatmasker.org*>. (2013-2015).
45. A. R. Quinlan, I. M. Hall, BEDTools: a flexible suite of utilities for comparing genomic features. *Bioinformatics* **26**, 841-842 (2010).
46. S. Koren *et al.*, Canu: scalable and accurate long-read assembly via adaptive k-mer weighting and repeat separation. *Genome Res* **27**, 722-736 (2017).
47. M. J. Chaisson, G. Tesler, Mapping single molecule sequencing reads using basic local alignment with successive refinement (BLASR): application and theory. *Bmc Bioinformatics* **13**, (2012).
48. P. Lindenbaum, JVarkit: java-based utilities for Bioinformatics. *figshare*, (2015).
49. I. Sovic *et al.*, Fast and sensitive mapping of nanopore sequencing reads with GraphMap. *Nat Commun* **7**, (2016).
50. M. Krzywinski *et al.*, Circos: An information aesthetic for comparative genomics. *Genome Research* **19**, 1639-1645 (2009).
51. B. Gel, E. Serra, karyoploteR: an R/Bioconductor package to plot customizable genomes displaying arbitrary data. *Bioinformatics* **33**, 3088-3090 (2017).

52. S. J. McIlwain *et al.*, Genome Sequence and Analysis of a Stress-Tolerant, Wild-Derived Strain of Saccharomyces cerevisiae Used in Biofuels Research. *G3-Genes Genom Genet* **6**, 1757-1766 (2016).
53. L. Zapata *et al.*, Chromosome-level assembly of Arabidopsis thaliana Ler reveals the extent of translocation and inversion polymorphisms. *P Natl Acad Sci USA* **113**, E4052-E4060 (2016).
54. K. E. Kim *et al.*, Long-read, whole-genome shotgun sequence data for five model organisms. *Sci Data* **1**, (2014).
55. R. Williams *et al.*, Amplification of complex gene libraries by emulsion PCR. *Nat Methods* **3**, 545-550 (2006).
56. E. Kalle, M. Kubista, C. Rensing, Multi-template polymerase chain reaction. *Biomol Detect Quantif* **2**, 11-29 (2014).
57. F. Madeira *et al.*, The EMBL-EBI search and sequence analysis tools APIs in 2019. *Nucleic Acids Res* **47**, W636-W641 (2019).
58. J. Hellemans, G. Mortier, A. De Paepe, F. Speleman, J. Vandesompele, qBase relative quantification framework and software for management and automated analysis of real-time quantitative PCR data. *Genome Biol* **8**, R19 (2007).
59. M. Serif *et al.*, One-step generation of multiple gene knock-outs in the diatom Phaeodactylum tricornutum by DNA-free genome editing. *Nat Commun* **9**, (2018).
60. M. A. Barbera, T. D. Petes, Selection and analysis of spontaneous reciprocal mitotic cross-overs in Saccharomyces cerevisiae. *Proceedings of the National Academy of Sciences of the United States of America* **103**, 12819-12824 (2006).
61. Y. Sui *et al.*, Genome-wide mapping of spontaneous genetic alterations in diploid yeast cells. *Proc Natl Acad Sci U S A*, (2020).
62. S. L. Amarasinghe *et al.*, Opportunities and challenges in long-read sequencing data analysis. *Genome Biol* **21**, (2020).
63. T. Rausch, M. H. Y. Fritz, J. O. Korbel, V. Benes, Alfred: interactive multi-sample BAM alignment statistics, feature counting and feature annotation for long- and short-read sequencing. *Bioinformatics* **35**, 2489-2491 (2019).
64. S. N. Lu *et al.*, CDD/SPARCLE: the conserved domain database in 2020. *Nucleic Acids Research* **48**, D265-D268 (2020).
